# Neuronal APOE4-induced Early Hippocampal Network Hyperexcitability in Alzheimer’s Disease Pathogenesis

**DOI:** 10.1101/2023.08.28.555153

**Authors:** Dennis R. Tabuena, Sung-Soo Jang, Brian Grone, Oscar Yip, Emily A. Aery Jones, Jessica Blumenfeld, Zherui Liang, Nicole Koutsodendris, Antara Rao, Leonardo Ding, Alex R. Zhang, Yanxia Hao, Qin Xu, Seo Yeon Yoon, Samuel De Leon, Yadong Huang, Misha Zilberter

## Abstract

The full impact of apolipoprotein E4 (APOE4), the strongest genetic risk factor for Alzheimer’s disease (AD), on neuronal and network function remains unclear. We found hippocampal region-specific network hyperexcitability in young APOE4 knock-in (E4-KI) mice which predicted cognitive deficits at old age. Network hyperexcitability in young E4-KI mice was mediated by hippocampal region-specific subpopulations of smaller and hyperexcitable neurons that were eliminated by selective removal of neuronal APOE4. Aged E4-KI mice exhibited hyperexcitable granule cells, a progressive inhibitory deficit, and E/I imbalance in the dentate gyrus, exacerbating hippocampal hyperexcitability. Single-nucleus RNA-sequencing revealed neuronal cell type-specific and age-dependent transcriptomic changes, including Nell2 overexpression in E4-KI mice. Reducing Nell2 expression in specific neuronal types of E4-KI mice with CRISPRi rescued their abnormal excitability phenotypes, implicating Nell2 overexpression as a cause of APOE4-induced hyperexcitability. These findings highlight the early transcriptomic and electrophysiological alterations underlying APOE4-induced hippocampal network dysfunction and its contribution to AD pathogenesis with aging.

---

The ε4 allele of apolipoprotein E (*APOE4*) is the strongest genetic risk factor for Alzheimer’s disease (AD) and lowers the age of onset of AD in a gene dose-dependent manner^1,2^. In most clinical studies, APOE4 carriers account for 60–75% of all AD cases^3^, highlighting the importance of APOE4 in AD pathogenesis. Longitudinal studies in humans demonstrate that APOE4’s detrimental effect on cognition is age-dependent and occurs before typical signs of AD arise^4^, with APOE4 carriers exhibiting age-related memory decline earlier in life than noncarriers^5^. Furthermore, structural imaging and cognitive studies have shown that healthy APOE4 carriers under the age of 40 exhibit reduced cortical thickness^6^ as well as gray matter atrophy and worse cognitive performance^7^. Human *APOE4* knock-in (E4-KI) mice lacking Aβ accumulation display sex- and age-dependent learning and memory deficits^8–11^. Together, available data suggest that APOE4 plays a role in AD not just in its active stages, but actually for decades before disease onset, including the preclinical and prodromal stages. Yet, there is still no clear understanding of how APOE4 expression promotes AD initiation and progression.

Early network hyperactivity in the form of subclinical seizures and interictal discharges is common in the preclinical and prodromal stages of AD and is associated with an accelerated cognitive decline^12^. Multiple transgenic AD mouse models exhibit network hyperexcitability and seizures preceding Aβ plaque formation^13–15^, and we have previously shown that epileptogenesis in APP/PS1 mice^16^ is driven by Aβ-induced neuronal hyperexcitability^15,17^. Hippocampal interictal spikes (IIS) in AD mice were shown to resemble those recorded in human epilepsy patients^18^. Importantly, this hallmark epileptiform activity inversely correlated with memory performance^18^, and other studies found that reducing hippocampal hyperexcitability with the antiepileptic drug levetiracetam improved cognition in amnesic mild cognitive impairment (MCI) patients^19,20^, as well as in APP/PS1^21^ and other AD mouse models^22,23^. All available data indicates that network hyperexcitability plays a role in AD-associated cognitive decline, positioning it as a potential target for effective treatment strategy^24^.

APOE4 is associated with region-specific hippocampal hyperactivity in non-demented young carriers as well as in MCI and early AD patients^25,26^, the extent of which predicts future memory decline^25^. Additionally, APOE4 has been linked to subclinical epileptiform activity during stress^24^. It is also a known risk factor for acquired epilepsy (AE), with APOE4 carriers at higher risk for temporal lobe epilepsy (TLE)^24,26^ and earlier TLE onset, especially after post-traumatic injury^27,28^. Moreover, APOE4 confers a six-fold higher probability of developing TLE treatment resistance^29^ and is associated with diminished memory performance in TLE patients^30,31^. Likewise, young E4-KI mice also spontaneously develop seizures starting at 5 months of age which are not observed in E3-KI or E2-KI animals, with females exhibiting higher seizure penetrance^32^. Several potential mechanisms responsible for early APOE4-related neuronal hyperactivity^33^ and network dysfunction have been reported, foremost among them inhibitory dysfunction^33^ and interneuron death^8,9,34 35–41^. However, the exact underlying neuronal mechanisms behind APOE4-induced network hyperexcitability remain unresolved. Interestingly, we have previously shown that reducing early network hyperactivity in 9 month-old E4-KI mice by 6-week pentobarbital treatment prevented inhibitory neuron death and cognitive deficits at 16 months^42^, suggesting that inhibitory dysfunction and learning impairment at more advanced ages may be a direct consequence of early network hyperactivity. Uncovering the causes of APOE4-driven early network pathophysiology is critical for advancing our understanding of the role APOE4 plays in AD pathogenesis.

In the current study, we set out to investigate the mechanisms and consequences of APOE-, age-, and region-dependent hippocampal network hyperexcitability. We used a multi-level analysis, including in vivo local field potential (LFP) data from freely moving mice^11^, whole-cell patch electrophysiology in ex vivo hippocampal slices, and single-nucleus RNA-sequencing (snRNA-seq), to identify the differentially expressed genes (DEGs) that may underly the neuronal functional phenotype across different ages. Finally, we validated select genes’ roles using targeted CRISPR interference (CRISPRi) in mediating APOE4-induced selective hippocampal neuronal hyperactivity.

Our findings demonstrate region-dependent hippocampal hyperexcitability in young E4-KI mice, which predicts learning and memory impairment in old age. This pathophysiology is driven by neuronal APOE4 expression, which elicits key transcriptomic alterations including Nell2 overexpression, resulting in neuronal hyperexcitability. The resulting region-specific network hyperactivity is further exacerbated by hilar inhibitory deficit and E/I imbalance with aging, all in concert contributing to APOE4-induced pathophysiology and cognitive impairments in an age-dependent manner.

## Results

### Region-specific hippocampal network hyperexcitability in E4-KI mice starting from young ages

Female E4-KI mice develop spontaneous seizures from the age of 5 months^32^, indicating that APOE4-induced pathophysiological changes in network excitability occur prior to the manifestation of observable cognitive deficits^9^. Although increased seizure susceptibility^43^ and phenotype^32^ in aged E4-KI mice have been reported, few studies examined the network mechanisms underlying these pathological phenotypes. While neuronal hyperactivity has previously been reported in the entorhinal cortex of aged E4-KI mice^33^, there is a lack of data on age-dependent APOE4 effects on network excitability in the hippocampus—a critical brain region for spatial learning and memory that is affected early in AD^44–46^. To explore age- and APOE4-dependent network activity parameters, we analyzed in vivo local field potential (LFP) data from our recent study on two independent cohorts of young (5–10 months) and aged (12–18 months) E-KI mice^11^. We focused on interictal spikes (IIS; Fig. 1a), a primary network hyperexcitability LFP biomarker and a hallmark of epilepsy observed in Alzheimer’s disease (AD) patients^12,47–49^ and most AD mouse models^13,16,17,50–52^. IIS represent spontaneous, high-amplitude, and brief synchronous discharges of a population of neurons^53^. We analyzed IIS rates in the cell layers of hippocampal CA1, CA3, and dentate gyrus (DG) regions during periods of activity. We found early APOE4-induced region-specific network hyperexcitability: starting from young ages, E4-KI mice exhibited increased IIS rates in CA3 (Fig. 1b) and DG (Fig. 1c). Meanwhile, we found no differences in IIS rates in CA1 (Fig. 1d), suggesting lack of network hyperexcitability in that region.

**Figure. 1.**
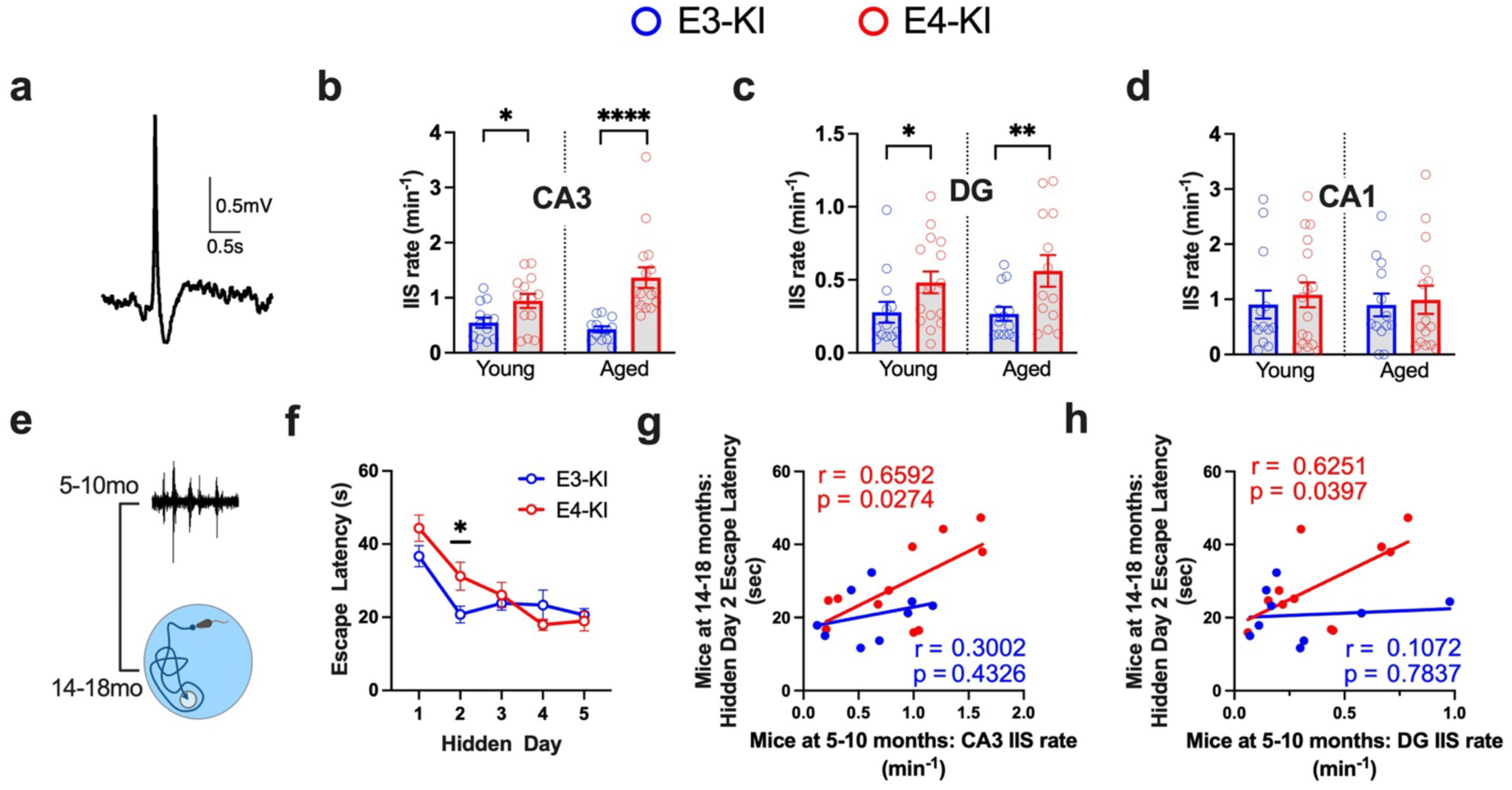
APOE4-driven hippocampal network hyperexcitability. **a,** Example raw LFP trace of an interictal spike from CA3 pyramidal cell layer. **b,** Interictal spike rates in CA3 of young (5–10) and aged (12–18) E3-KI and E4-KI mice. **c,** Dentate spike rates in DG of young and aged E3-KI and E4-KI mice. **d,** Interictal spike rates in CA1 of young and aged E3-KI and E4-KI mice. **e,** Timeline of experiments for the longitudinal cohort shown in (f) and (g). **f,** Average daily escape latency on MWM for the longitudinal E3-KI and E4-KI cohort at 14–18 month of age. **g,** CA3 interictal spike rate at 5–10 months predicts hidden day 2 escape latency at 14–18 months in E4-KI but not in E3-KI mice. Pearson’s correlation analysis (two-sided). **h,** DG interictal spike rate at 5–10 months predicts hidden day 2 escape latency at 14–18 months in E4-KI but not in E3-KI mice. Pearson’s correlation analysis (two-sided). All data are represented as mean ± s.e.m., *p<0.05, **p<0.01, ****p<0.0001, unpaired two-tailed t-test or Mann-Whitney U test. For IIS analysis, N=14 and 15 for young E3-KI and E4-KI mice; N=13 and 16 for aged E3-KI and E4-KI mice. MWM test, N=9 and 12 for E3-KI and E4-KI mice.

Since young E4-KI mice showed no detectable spatial learning deficit in a Morris water maze (MWM) test^11^, we examined whether early network hyperexcitability in young E4-KI mice is predictive of future behavioral deficits in a longitudinal cohort, in which the LFP recordings were conducted at young (5–10 months) ages and spatial learning and memory tests were done at old (14–18 months) ages (Fig. 2e). As expected, aged (14–18 months) E4-KI mice exhibited impaired spatial learning in the MWM test, seen as increased escape latency on hidden day 2 (Fig. 1f). Strikingly, both CA3 and DG IIS rates in young E4-KI mice (5–10 months) showed a significant correlation with hidden day 2 escape latency at old ages (14–18 months; Fig. 1g,h), indicating that network hyperexcitability at young ages predicts spatial learning deficits at old ages. In contrast, neither CA3 nor DG IIS rates in young E3-KI mice correlated with hidden day 2 escape latency at old ages (Fig. 1g,h). These results demonstrate that APOE4-induced early hippocampal network hyperexcitability, especially in the CA3 and DG subregions, represents a major “preclinical” pathophysiology predicting cognitive impairment at old ages.

**Figure. 2.**
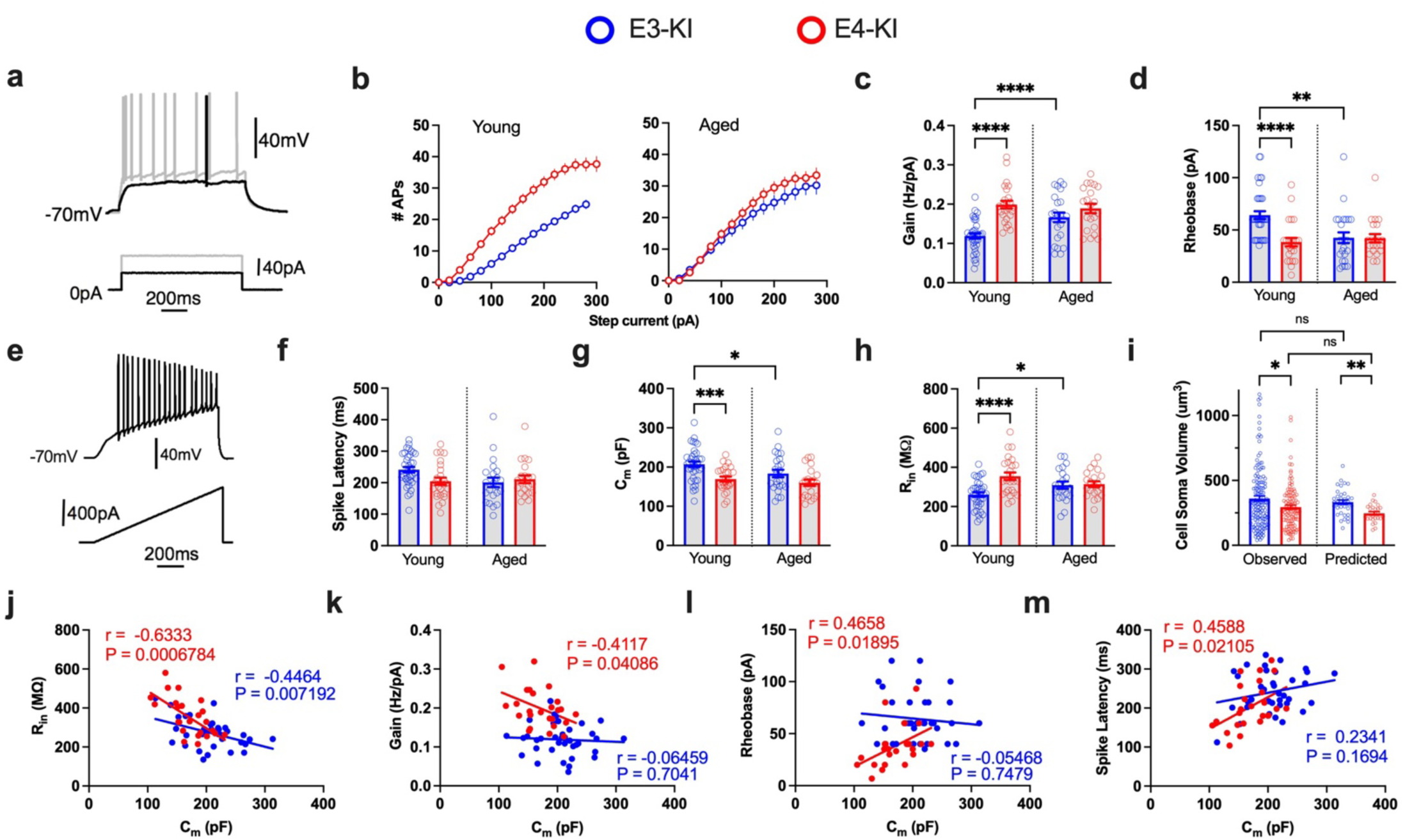
APOE4-driven intrinsic electrophysiological differences in hippocampal CA3 pyramidal cells (PCs). **a,** Example whole-cell current clamp recording of a CA3 PC with membrane potential response (top) to a 1-s depolarizing current injection (bottom) at rheobase (threshold current; black) and 2x rheobase (grey). **b,** Input current – firing rate relationship (I-F Curve), the average number of spikes elicited by 1sec epochs of incremental 20pA depolarizing current injection. **c,** Firing gain values, quantified as the linear slope of the I-F curve. **d,** Rheobase values, representing the minimum 1s depolarizing current required to elicit an action potential. **e,** Example whole-cell current clamp recording of spike latency protocol with membrane potential (top) response to a ramped 800pA, 1-s depolarizing current injection (bottom)**. f,** Spike latency values. **g,** Cell membrane capacitance (C_m_) values. **h,** Input resistance (R_in_) values. **i,** Cell soma volumes for CA3 PCs, measured directly or predicted using reported specific C_m_ of 0.9pF/cm^2^. Statistical differences analyzed using Kolmogorov-Smirnov test. **j,** Correlation between C_m_ and R_in_ values in young E3-KI (blue) and E4-KI (red) CA3 PCs. Numbers represent Pearson’s correlation coefficient and corresponding p-values. **k–m,** Cell membrane capacitance correlates with neuronal excitability in E4-KI but not in E3-KI neurons: correlation between C_m_ and firing gain (k), rheobase (l), and spike latency (m). Pearson’s correlation analysis (two-sided). All data are represented as mean ± s.e.m., *p<0.05, **p<0.01, ***p<0.001, ****p<0.0001, unpaired two-tailed t-test or Mann-Whitney U test. N= 8 and 7 for young E3-KI and E4-KI mice, respectively; N= 6 and 4 for aged E3-KI and E4-KI mice

### APOE4 induces cell type-specific neuronal hyperexcitability in the hippocampus of young mice

We next investigated the neuronal mechanisms underlying the region-specific network hyperexcitability in the hippocampus of E4-KI mice. Neuronal intrinsic excitability (IE) is represented by a number of fundamental parameters typically recorded via whole-cell patch clamp. We restricted our focus to rheobase (minimal depolarizing current required for activation; Fig. 2a), firing gain (relationship between incremental 1-s depolarizing current step amplitude and the resulting spiking frequency), and spike latency (delay to first action potential in response to a 1-s 800pA ramped current injection; Fig. 2e). We also examined passive neuronal properties modulating excitability, such as input resistance (R_in_) as well as cell membrane capacitance (C_m_) which is proportional to the membrane surface area and reflects neuronal size^54^.

#### CA3 pyramidal cells

In young E4-KI mice (7–9 months of age), CA3 pyramidal cells (PCs) exhibited pronounced hyperexcitability compared to neurons from young E3-KI mice (Figure 2). CA3 PCs had higher firing gain (Fig. 2b,c) as well as lower rheobase values (Fig. 2d), while APOE genotype-mediated differences in spike latency were not significantly altered (Fig. 2e,f). Interestingly, in aged E3-KI mice (17-19 months of age), CA3 PCs showed significantly increased excitability (Fig. 2b–f) compared to young E3-KI animals, and were similar to neurons from aged E4-KI mice, eliminating the difference in excitability between old E4-KI and E3-KI mice. We also compared other IE-related parameters, such as the resting membrane potential (RMP), maximal firing rate, and spike accommodation (Extended Data Fig. 1), as well as action potential (AP) waveform parameters (Extended Data Fig. 2). We found elevated maximal firing rate (Extended Data Fig. 1b) and increased fast afterhyperpolarization (fAHP) (Extended Data Fig. 2e) in young E4-KI versus E3-KI mice, and no differences in AP amplitude, rise time, half-width, or slow afterhyperpolarization (sAHP) (Extended Data Fig. 2a–d,f). CA3 PCs in young E4-KI mice also had smaller C_m_ values (Fig. 2g) paralleled by higher input resistance (Fig. 2h,i). Since specific membrane capacitance in neurons is constant at 0.9μF/cm^2^ across all cell types^55^, C_m_ is a direct representation of total membrane surface area and therefore, cell size^54^. We validated this relationship by directly measuring cell soma volume of CA3 PCs from young E3-KI and E4-KI mice using laser scanning confocal microscopic images, and found that observed cell volumes were remarkably similar to those predicted by the C_m_ measurement (Fig. 2i). Observed CA3 PC volumes also confirmed that cells from young E4-KI mice were smaller compared to those from young E3-KI mice (Fig. 2i). Furthermore, C_m_ and R_in_ values were significantly correlated in both E3-KI and E4-KI neurons (Fig. 2j), verifying that cell size mediates cell input resistance, as reported previously^56,57^. Similar to age-related IE changes in E3-KI mice, CA3 PCs from aged E3-KI mice became smaller as evidenced by significantly lower C_m_ and higher R_in_ values compared to those in young E3-KI mice (Fig. 2g,h). The apparent similarity between age- and genotype-dependent changes in cell size and IE suggests that these neuronal properties might be related, as shown previously in other AD models^58,59^. To uncover this relationship, we explored the correlations between C_m_ and primary IE parameters in CA3 PCs from young E-KI mice. In E4-KI CA3 PCs, every IE parameter correlated significantly with C_m_, while the corresponding parameter in E3-KI cells showed no such correlation (Fig. 2k–m), indicating that APOE4-induced hyperexcitability is indeed, at least in part, underlain by reduced cell size.

#### Dentate granule cells

Among recorded dentate granule cells (DGCs), we found two distinct populations based on their electrophysiological properties. Type I DGCs^60^ were characterized by their inability to sustain firing under higher stimulation intensities (Fig. 3a,b) and displayed a smaller cell size (lower C_m_ and higher R_in_ values). This was paralleled by a hyperexcitability phenotype in both E3-KI and E4-KI mice at both ages, as indicated by higher firing gain, lower rheobase, and lower spike latency compared to Type II DGCs^60^ (Extended Data Fig. 3a–j) which displayed sustained firing under higher stimuli (Fig. 3a, b). While the relative population of Type I DGCs did not differ between young E3-KI and E4-KI mice (33% in E3-KI vs. 35% in E4-KI) (Fig. 3c), aged E4-KI mice possessed a significantly higher proportion of Type I DGCs (43% in E3-KI vs. 64% in E4-KI; p = 0.0008, Cochran-Mantel-Haenszel test). Interestingly, the cell size and hyperexcitability phenotypes in E4-KI mice were restricted to young Type II DGCs, which exhibited smaller C_m_ (Fig. 3i) and higher R_in_ (Fig. 3j), as well as lower rheobase (Fig. 3l) and spike latency (Fig. 3m) values. Confirming that reduced cell size mediates hyperexcitability in young E4-KI Type II DGCs, C_m_ values correlated with rheobase and spike latency values in E4-KI but not E3-KI neurons (Extended Data Fig. 3l–n). Finally, we found no APOE4-mediated size or excitability differences in Type II DGCs of aged E-KI mice, again due to E3-KI cells becoming smaller and more excitable in aged mice (Fig. 3i,j,m). However, as Type I DGCs were hyperexcitable compared to Type II DGCs (Fig. 3b), their increased fraction likely contributed to a persistent combined DGC hyperexcitability phenotype in aged E4-KI mice (Fig. 3n–r).

**Figure 3.**
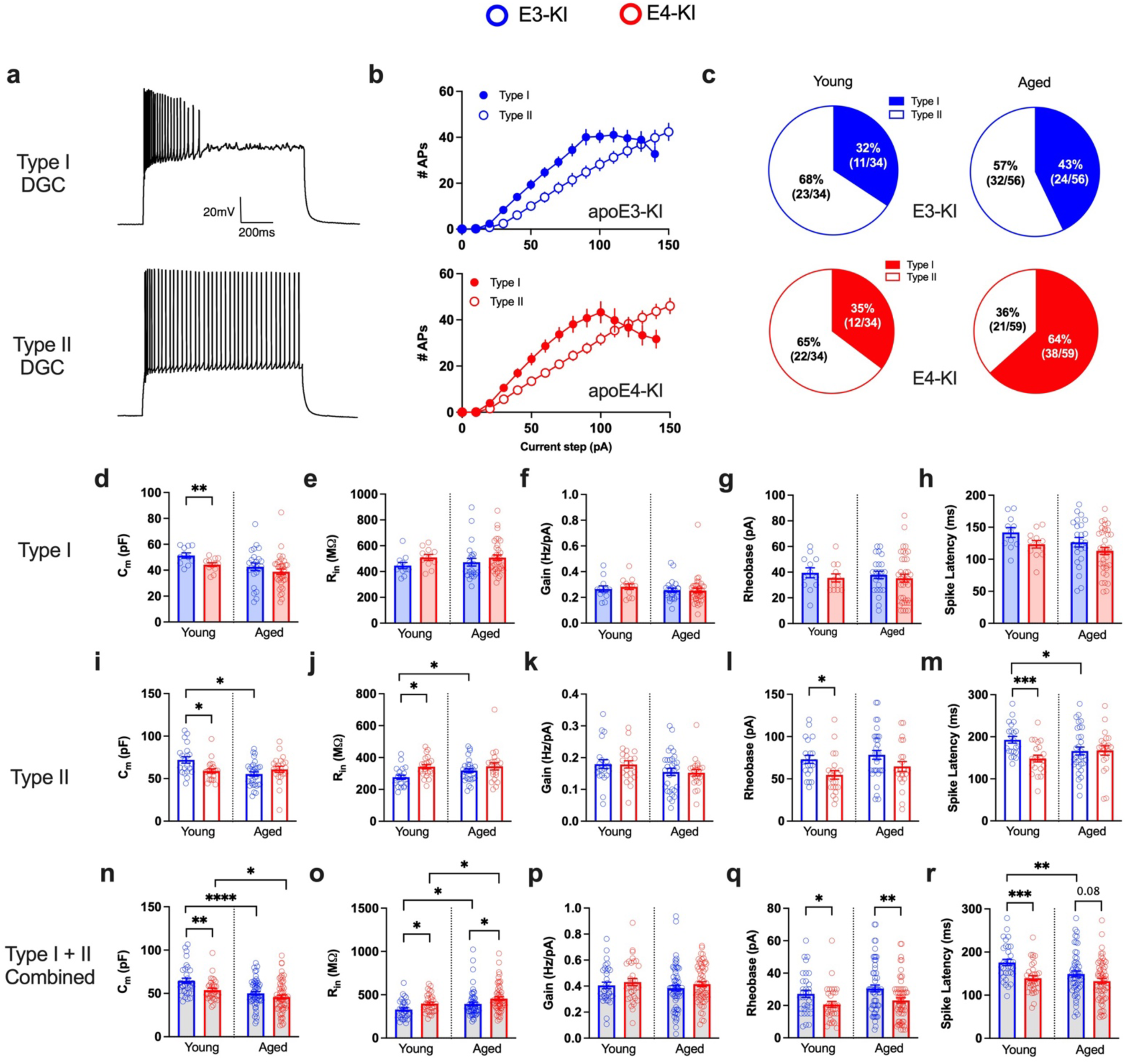
Dentate Gyrus granule cell (DGC) subpopulations display age-, APOE genotype- and cell subtype-specific morpho-electric and excitability differences. **a,** Representative DGC firing traces in response to a 300pA 1-s depolarizing current injection. **b,** Input current – firing rate relationships (I-F Curve) in DGCs from young E3-KI (blue) and E4-KI (red) mice: the number of spikes elicited by 1 second epochs of incremental 10pA depolarizing current injections. **c,** Proportions of Type I DGCs and Type II DGCs in E3-KI (blue) and E4-KI (red) young and aged mice. **d–h,** Lack of age- or APOE-related IE phenotype in Type I DGCs: values for C_m_ (d), R_in_ (e), Firing gain (f), Rheobase (g), and spike latency (h). **i–m,** Age- and APOE-dependent changes in morpho-electric parameters of Type II DGCs: values for C_m_ (i), R_in_ (j), Firing gain (k), Rheobase (l), and spike latency (m). **n–r,** Age- and APOE-dependent changes in morpho-electric parameters in combined DGC population: values for C_m_ (n), R_in_ (o), Firing gain (p), Rheobase (q), and spike latency (r). All data are represented as mean ± s.e.m., *p<0.05, **p<0.01, ***p<0.001, ****p<0.0001, unpaired two-tailed t-test or Mann-Whitney U test. N= 4 and 3 for young E3-KI and E4-KI mice; N= 5 each for aged E3-KI and E4-KI mice.

#### CA1 PCs

In line with the observed lack of APOE4-induced network excitability phenotype in CA1 (Fig. 1d), we found no significant IE phenotype in CA1 PCs of E4-KI mice versus those of E3-KI mice at both young and old ages (Extended Data Fig. 4b–f), indicating neuronal cell type specificity of the APOE4 effect. Although CA1 PCs from E4-KI mice did exhibit smaller C_m_ values in both age groups compared to those from E3-KI mice (Extended Data Fig. 4a), the difference was less pronounced (C_m_ in young E4-KI mice: CA1 PCs 87%, CA3 PCs 82%, Type II DGCs 82% vs. young E3-KI mice) and did not translate to significant differences in R_in_ (Extended Data Fig. 4b). We also found no differences in secondary excitability parameters of CA1 PCs across APOE genotypes (Extended Data Figs. 2g–i and 3m–r).

Taken together, young E4-KI mice exhibited a cell type-specific hyperexcitability phenotype in CA3 PCs and DGCs which was associated with reduced cell size. Aging did not affect this phenotype in E4-KI mice, but did result in a significantly increased population of hyperexcitable Type I DGCs, promoting dentate hyperexcitability in aged E4-KI mice. In contrast, we found smaller and more excitable CA3 PCs and DGCs in aged E3-KI mice compared to young mice, indicating an age-dependent effect of APOE3 toward APOE4’s detrimental influence that starts early in life.

### Removal of APOE4 from neurons rescues the hyperexcitability phenotype in young E4-KI mice

APOE is largely made by astrocytes in the brain, although it can also be produced in stressed neurons^61–63^. To find out whether APOE4-related morpho-electric and excitability phenotypes were caused by neuronal or astrocytic APOE4 expression^64,65^, we performed similar experiments in young E4-KI mice in which the *APOE4* gene was selectively deleted in either neurons (fE4-KI^Syn1-Cre^ mice^64^) or astrocytes (fE4-KI^GFAP-Cre^ mice^64^). By recording passive and active properties of CA3 PCs from these mice, we observed that neuronal APOE4 removal resulted in full normalization of all altered parameters — C_m_ and R_in_, firing gain, rheobase, and spike latency — to the levels seen in E3-KI mice (Fig. 4a–f). This highlights the crucial role of neuronal APOE4 in inducing morpho-electric and excitability changes. Furthermore, deleting neuronal APOE4 eliminated all correlations between C_m_ and primary IE parameters (Fig. 4g–i), supporting the link between neuronal APOE4-induced cell atrophy and hyperexcitability. In contrast, removing astrocytic APOE4 did not significantly affect electrophysiological properties reflecting neuronal cell size (Extended Data Fig. 5a,b) or excitability (Extended Data Fig. 5c–f). These findings underscore the pivotal role of neuronal APOE4 in driving neuronal atrophy and consequent hyperexcitability.

**Figure 4.**
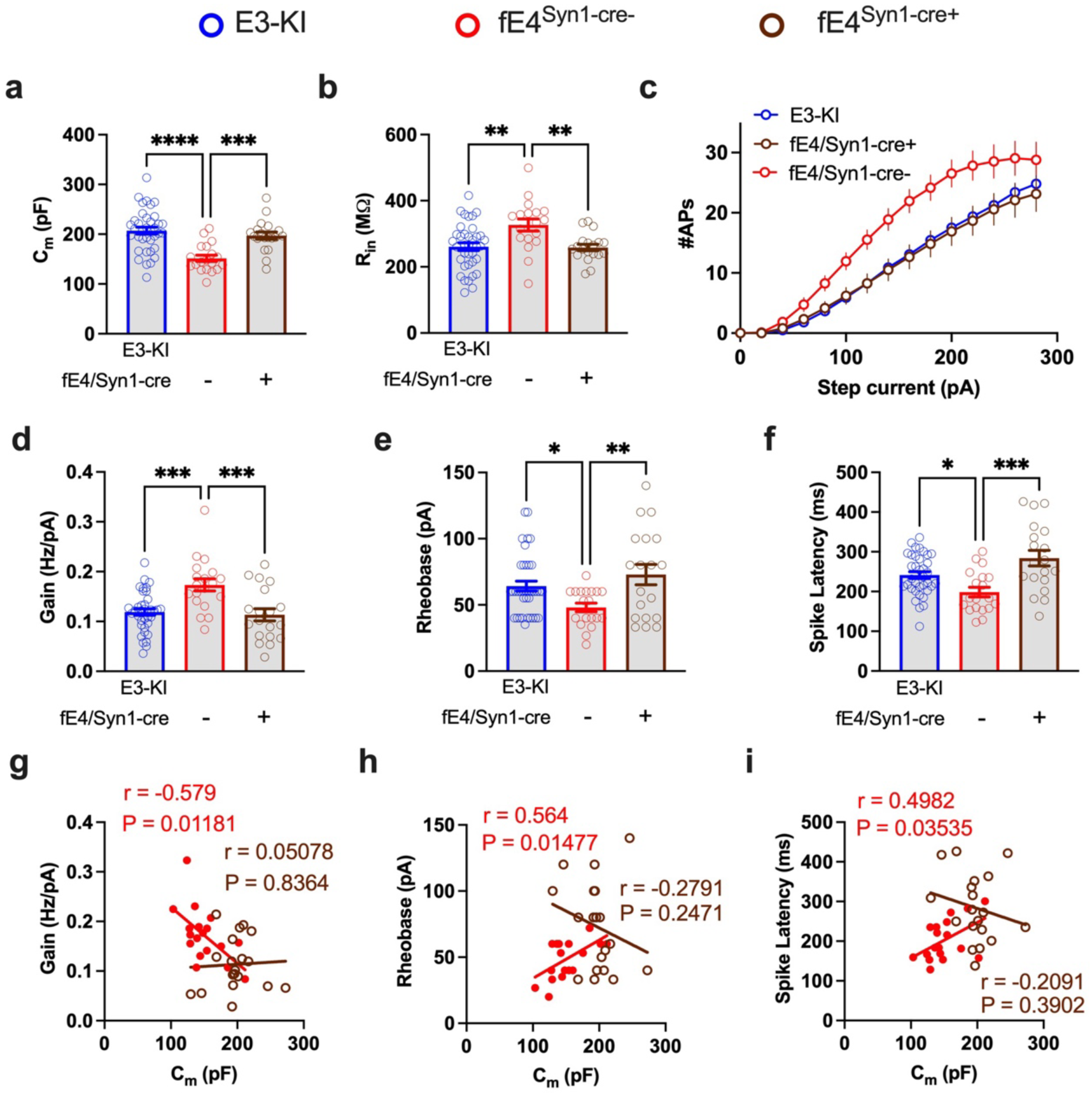
Selective APOE4 removal from neurons rescues morpho-electric and excitability phenotypes of CA3 PCs in young fE4-KI^Syn1-Cre+^ mice. Compared to their Cre-negative littermates that express APOE4 in neurons (red), young fE4-KI^Syn1-Cre+^ mice (brown) display full normalization of all morpho-electric and excitability parameters to E3-KI levels (blue). **a,** C_m_ values. **b,** R_in_ values. **c,** I-F curves. **d,** Firing gain values. **e,** Rheobase values. **f,** Spike latency values. **g–h,** Neurons in fE4-KI^Syn^^1-Cre+^ mice exhibit abolished C_m_ - IE correlations, while neurons in Cre-negative mice maintain these correlations in C_m_ vs. Firing gain (g), C_m_ vs Rheobase (h), and C_m_ vs. spike latency (i). All data are represented as mean ± s.e.m., *p<0.05, **p<0.01,***p<0.001,****p<0.0001, one-way ANOVA with Tukey’s post-hoc multiple comparisons test. N= 8 for E3-KI mice and 3 for each fE4-KI^Syn1-Cre^ genotype.

## Selective vulnerability of some of the CA3 PCs and DGCs to APOE4-induced hyperexcitability

Significant within-group heterogeneity and overlap in the neural parameters between genotypes (Figs. 2,3) suggest the presence of distinct sub-populations of pathological and healthy cells among CA3 PCs and DGCs. To identify such, we employed the k-means method to cluster neurons based on their morpho-electric properties. We pooled CA3 PCs from all genotypes and ages, normalizing values prior to clustering (Fig. 5a). We then used k-means clustering to identify two clusters (*k = 2*), with each cluster described by its *centroid*, the set of mean parameter values (Fig. 5b). Clusters clearly divided primarily along morpho-electric and excitability parameters that were significantly different between young E4-KI and E3-KI CA3 PCs and not along parameters that did not show differences across APOE genotypes and ages (Fig. 5b, Fig. 2, and Extended Data Figs. 2 and 3), indicating that the clustering analysis captured key differences between pathologically hyperexcitable and healthy or normal neurons. To validate the results and ensure the robustness of the centroid characteristics, we utilized a five-fold cross-validation (see methods) to determine that two clusters (Fig. 5c) and not three (Fig. 5d) yielded robust clustering with stable centroids across 1000 iterations. This strongly suggests only two primary sub-populations of neurons divided by key morpho-electric parameters (Fig. 5b).

**Figure 5.**
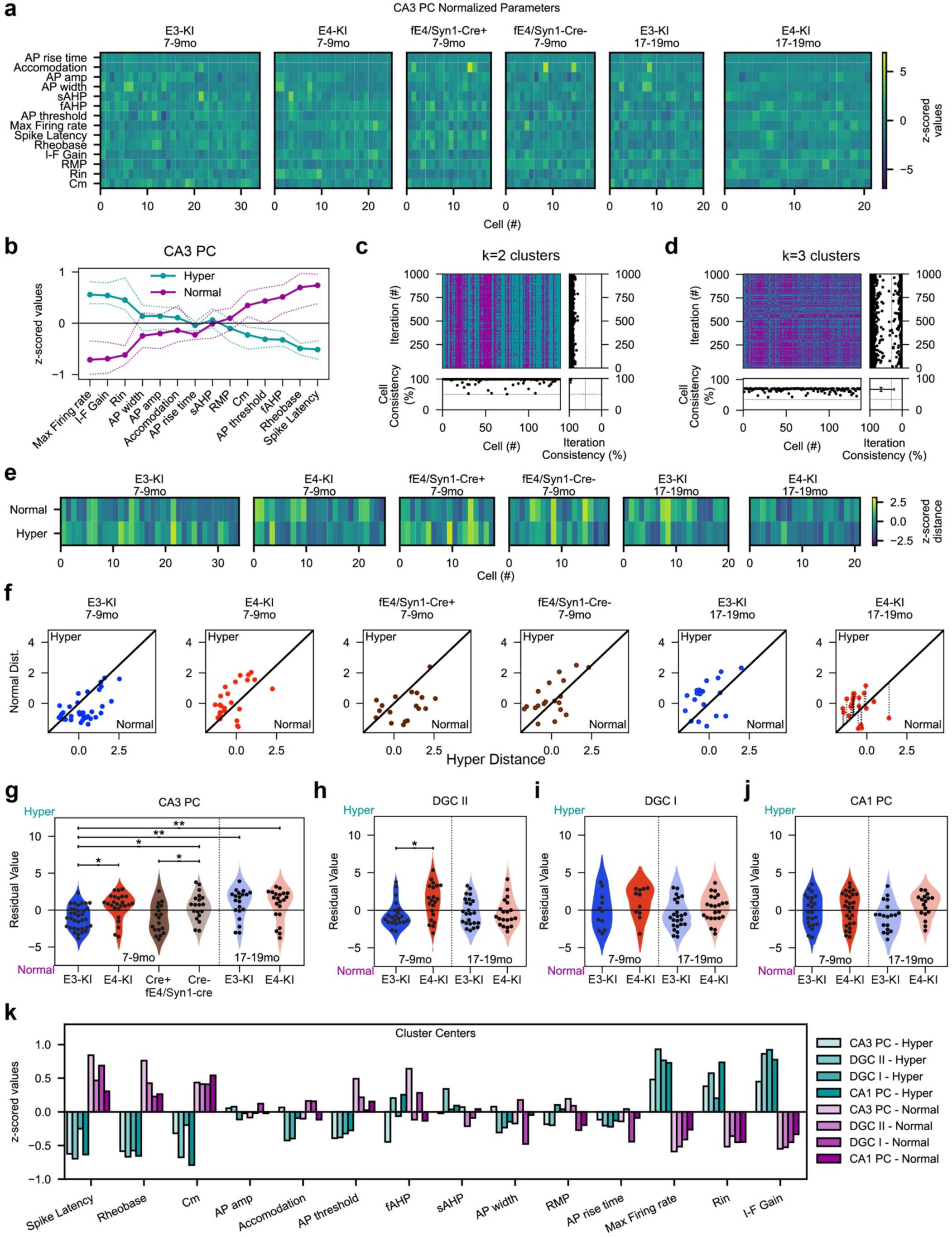
Cluster analysis with k-means reveals selective vulnerability of CA3 PCs and DGCs. **a,** Normalized cell parameters for CA3 PCs used for clustering analysis. **b,** Centroids from k-means with 2 clusters. Solid lines indicate the mean centroid after 1000 cross-validation iterations with dashed lines indicating the boot strapped 99% confidence interval. Cells divide into normal and hyperexcitable subtypes. Morpho-electric parameters C_m_ and R_in_, strongly associate with increased active metrics of excitability (e.g., high firing rate gain and low rheobase). **c,** Cross validation of k-means sorting with k=2 clusters. Top left: Heat map showing the cluster label for each cell across all iterations. Top right: Raster showing consistency of labeling between full data set cluster and each iteration. Bottom left: Raster plot depicting consistency of labeling for each cell. Bottom right: Combined mean and standard deviation of cell and iteration consistency. **d,** Cross-validation as in (c) but with k=3 clusters. **e,** Distance from Normal and Hyperexcitable centroids for each cell group plotted as a heat map. **f,** Scatterplots for each cell depicting the normalized k-means distance from Normal and Hyperexcitable centroids. The decision line for clustering is the unity line plotted for each scatter plot. Right panel: the residuals for each cell are shown as dashed lines. Large positive residuals are more strongly associated with Hyperexcitable parameters while large negative residuals with Normal. **g–j.** Violin plots of the residual values for CA3 PCs (g), CA1 PCs (h), Type I DGCs (i), and Type II DGCs (j) across genotypes and age groups. **k,** Centroid loadings for all cell types. *p<0.05, **p<0.01, two-way ANOVA with post-hoc Tukey test.

We then examined the distance of each neuron from the cluster centroids (Fig. 5e), which reflects the similarity (small distance) or dissimilarity (large distance) of each cell to the cluster centroid. By plotting the cells’ distances on a scatter plot, the k-means classification could be visualized, with the decision (unity) line representing equidistance from each centroid (Fig. 5f). Each neuron’s signed distance to the unity line (its residual value; Figure 5f, last panel) is therefore representative of its strength of cluster membership. Analysis of membership strength revealed the impact of neuronal APOE4 expression and age on cell classification, with significantly more hyperexcitable neurons in young E4-KI mice compared to age-matched E3-KI mice, and more E3-KI neurons shifting into the hyperexcitable cluster with age (Fig. 5g). However, a proportion of E4-KI neurons did exhibit strong membership in the normal cluster at both ages, indicating selective vulnerability of CA3 PCs to APOE4 effects. Finally, deletion of neuronal APOE4 in young fE4-KI^Syn1-Cre+^ mice resulted in a significant depletion of the hyperexcitable CA3 PC cluster compared to their Cre-negative littermates, which resembled the E4-KI mouse cluster distribution (Fig. 5g).

The same analysis was performed for DGCs and CA1 PCs. Type II DGCs displayed similar patterns of hyperexcitability classification, primarily determined by differences in morpho-electric parameters R_in_ and C_m_ (Extended Data Fig. 6a). Significantly more type II DGCs in young E4-KI mice belonged to the hyperexcitable cluster compared to young E3-KI type II DGCs (Fig. 5h), while no age- or APOE genotype-dependent differences were observed in type I DGCs (Fig. 5i). Meanwhile, in CA1 PCs, no significant APOE genotype- or age-related differences in hyperexcitable populations were observed (Fig. 5j), despite robust clustering of hyperexcitable and normal cells (Extended Data Fig. 6c,f)

Comparing the centroids of all cell types highlighted consistent divisions, with hyperexcitable cells defined by morpho-electric parameters (R_in_ and C_m_) and IE parameters (firing gain, rheobase, and spike latency), but not by AP waveform or other membrane properties (Fig. 5k). Cluster enrichment was ultimately determined by cell type-, APOE genotype-, and age-mediated differences in these parameters, reinforcing our conclusion that neuronal APOE4 expression in select sub-populations of neurons results in reduced cell size and, consequently, intrinsic hyperexcitability.

### Region-specific and age-dependent synaptic excitation-inhibition imbalance in the hippocampus of E4-KI mice

Another major mechanism contributing to hyperactivity and cognitive deficits in E4-KI mice is synaptic dysfunction, especially the loss of inhibitory function^8,33^. To investigate the network and synaptic consequences of APOE4 expression, we recorded spontaneous excitatory post-synaptic currents (sEPSCs) and inhibitory post-synaptic currents (sIPSCs) in CA3 PCs and DGCs in both young and old age groups (Fig. 6). CA3 PCs in young E4-KI mice displayed higher sEPSC frequency and similar amplitude compared to those in young E3-KI mice (Fig. 6a,b and Extended Data Fig. 7a), consistent with hyperexcitability of these cells given the recurrent excitation in CA3 local networks^66^. We found no APOE genotype-specific differences in sIPSC frequency or amplitude in CA3 PCs between young E4-KI and E3-KI mice, indicating unaltered inhibitory tone across the APOE genotypes at this age (Fig. 6c,d; Extended Data Fig. 7c,d). Increased excitatory drive together with unchanged inhibitory input resulted in a higher excitatory-to-inhibitory (E-I) ratio in CA3 PCs in young E4-KI versus E3-KI mice (Fig. 6e). At old age, both the excitatory input frequency (Fig. 6a) and E-I ratio (Fig. 6e) significantly increased in CA3 PCs in E3-KI mice, abolishing the differences between APOE genotypes and mirroring the age-related trend observed in excitability of these cells.

**Figure 6.**
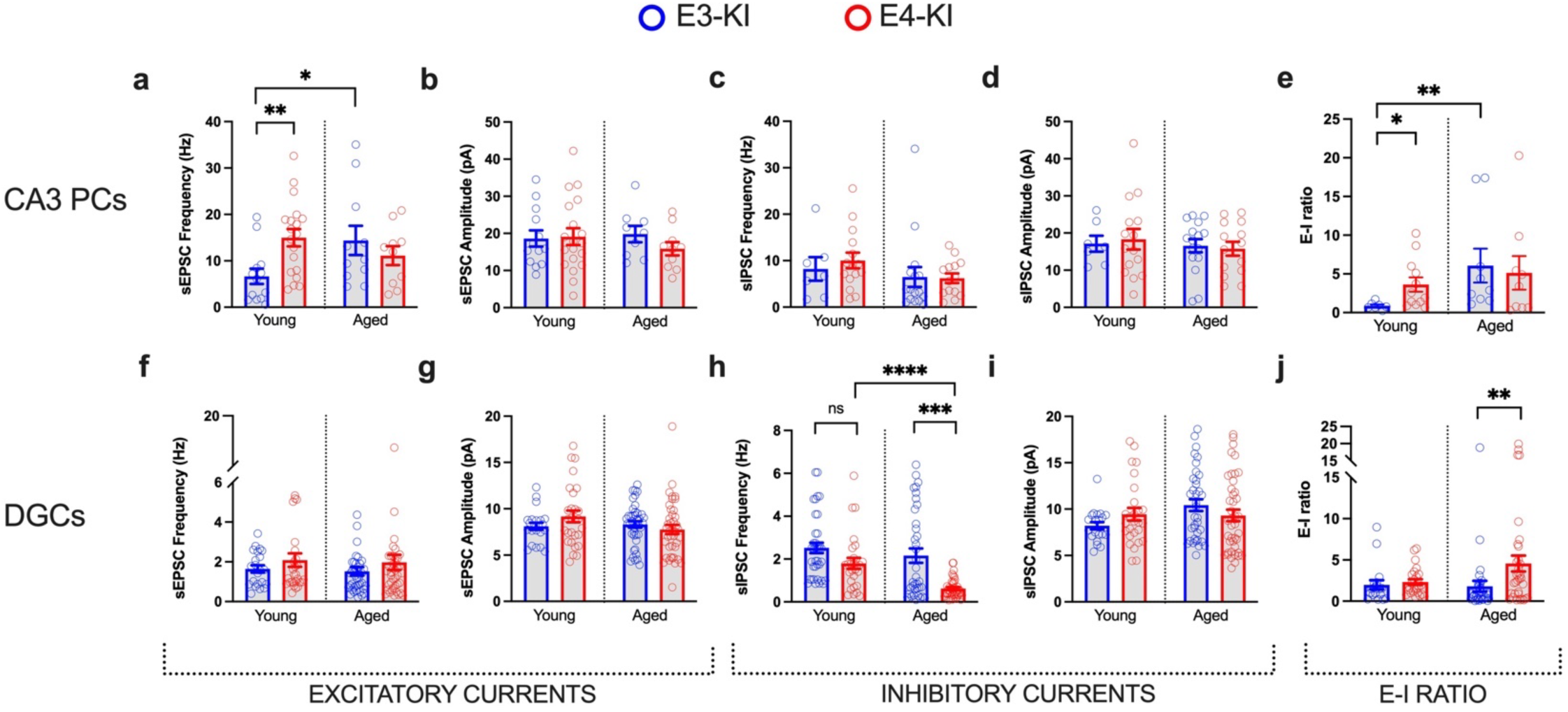
APOE4-induced changes in spontaneous synaptic activity in CA3 and DG in young and aged mice. **a,** Spontaneous excitatory post-synaptic currents (sEPSC) frequency in CA3 PCs. **b,** sEPSC amplitudes in CA3 PCs. **c,** Spontaneous inhibitory post-synaptic currents (sIPSC) frequency in CA3 PCs. **d,** sIPSC amplitudes in CA3 PCs. **e,** Excitation-Inhibition (E-I) ratio in CA3 PCs. **f,** sEPSC frequency in DGCs. **g,** sEPSC amplitudes in DGCs. **h,** sIPSC frequency in DGCs. **i,** sIPSC amplitudes in DGCs. **j,** E-I ratio in DGCs. All data are represented as mean ± s.e.m., *p<0.05; **p<0.01, ***p<0.001, ****p<0.0001, unpaired two-tailed t-test or Mann-Whitney test. N= 5 and 6 mice for CA3 PCs from young E3-KI and E4-KI mice, respectively; N= 6 for DGCs from young E3-KI and E4-KI mice. N= 6 and 5 mice for CA3 PCs from aged E3-KI and E4-KI mice; N= 7 and 10 for DGCs from aged E3-KI and E4-KI mice.

We found no differences in sEPSCs in DGCs from E4-KI mice at either age (Figs. 6f,g; Extended Data Fig. 7e,f). At the same time, in line with our previous reports on accelerating hilar interneuron death in E4-KI mice during aging^8,9^, DGCs in aged E4-KI mice displayed a progressive reduction in sIPSC frequency with age (Fig. 6h and Extended Data Fig. 7g), resulting in significantly higher E-I ratio (Fig. 6j) in old E4-KI versus E3-KI mice.

### Neuronal APOE4 expression leads to neuronal cell type-specific and age-dependent transcriptomic changes

The apparent heterogeneity of APOE4 effect in neuronal subtypes appears similar to the cell type selectivity of APOE4 effect on neuronal transcriptomes that we reported recently^62^, and suggests a potential transcriptomic signature underlies susceptibility and resilience of APOE4-expressing neurons. To uncover APOE4-related gene expression differences in different types of hippocampal neurons, we analyzed our published snRNA-seq datasets obtained from E3-KI and E4-KI mouse hippocampi at different ages (Fig. 7a,b)^62^. To explore the transcriptomic alterations associated with the protective effects of APOE4 deletion in neurons of young E4-KI mice (Fig. 4), we also generated snRNA-seq data from the hippocampi of 5- and 10-month-old female fE4-KI^Syn1-Cre+^ mice (Extended Data Fig. 8). To select differentially expressed genes (DEGs) whose transcriptional profile changes fit the observed neuronal morpho-electric phenotypes in an APOE genotype-, cell type-, and age-dependent manner, we focused on clusters 1 and 2 (DGCs) as well as 6 (CA2/3 PCs). As 5- and 10-month-old fE4-KI^Syn1-Cre+^ sequencing was done independently from our previous report^62^, the sequence data was clustered separately, aligning cluster identities with the original dataset clusters using the same marker genes, with cluster 1 aligned to original cluster 1, cluster 2 to 2, and cluster 11 to 6 (Extended Data Fig. 8, Supplementary Table 1). For further analysis, we looked for differentially expressed genes according to five selection criteria (Fig. 7c): 1) Candidate genes should be differentially expressed between E4-KI and E3-KI mice at 5 and 10 months of age. 2) Increased gene expression in cells with high APOE expression from 10mo E4-KI mice; we classified cells as APOE-expression-high if they express APOE mRNA at >2 SD above the median expression for that cell type. 3) Not differentially expressed in neurons from fE4KI^Syn1-Cre+^ mice versus those from E4-KI mice at 5 months or 10 months of age. 4) Not differentially expressed in in neurons from 20 month-old E4-KI mice versus 20-month-old E3-KI mice. 5) Finally, genes should be differentially expressed in neurons from 20-month-old E3-KI mice compared to neurons from 5-month-old E3-KI mice, with same directional change as in (1–2). This last criterion was important to include since we observed a significant reduction in C_m_ together with increased excitability of both CA3 PCs and DGCs in aged versus young E3-KI mice (Figs. 2,3). From the initial pool of 552 DEGs for DG cluster 1 (Supplementary Table 2), 889 for DG cluster 2 (Supplementary Table 3), and 492 for CA2/3 PC cluster 6 (Supplementary Table 4) in 5-month-old mice (Fig. 7c), our analysis pipeline yielded 2 final candidate DEGs for DG cluster 1, 13 for DG cluster 2, and 7 for CA2/3 PC cluster 6 (Fig. 7c, Supplementary Table 6). In contrast, the same DEG analysis on dorsal CA1 PCs (E-KI cluster 3 and fE4-KI^Syn1-cre+^ cluster 5; Supplementary Table 5) yielded no candidates, consistent with these neurons showing no APOE-dependent IE phenotype (Extended Data Fig. 4).

**Figure 7.**
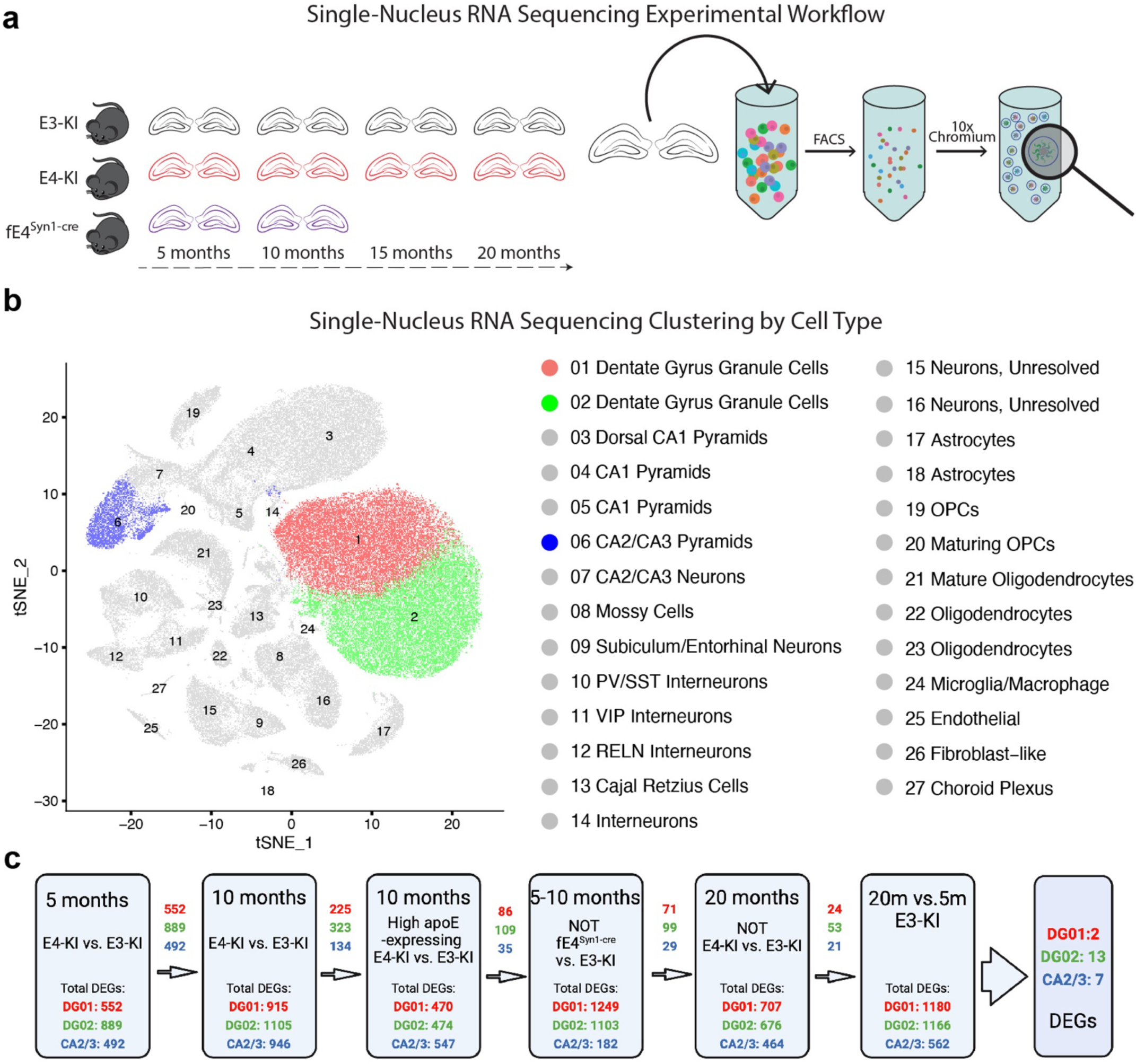
snRNA-seq analysis of E3-KI and E4-KI mouse hippocampi at different ages. **a,** Experimental design: hippocampi were extracted from female E3-KI and E4-KI mice at 5, 10, 15 and 20 months of age (N=4 per genotype and age), and from female fE4-KI^Syn1-cre+^ mice at 5 and 10 months of age. The hippocampi were dissociated, and nuclei were labeled with DAPI and isolated using flow cytometry before processing using the 10x Chromium v2 system for snRNA-seq. **b,** Clustering using the Seurat package revealed 27 distinct cellular populations in E3-KI and E4-KI mice. Marker gene analysis led to the identification of 16 neuronal clusters (Clusters 1–16) and 11 non-neuronal clusters (Clusters 17–27). **c,** DEG selection criteria for clusters 1, 2, and 6. Values represent the number of differentially expressed genes passing each successive criteria. See also Extended Data Fig. 8 and Supplementary Tables 1–5.

### Targeted reversal of Nell2 overexpression in neurons of young E4-KI mice rescues their smaller size and hyperexcitability phenotypes

Following the identification of a number of DEGs that may modulate the detrimental impact of APOE4 on neuronal excitability and/or susceptibility or resistance to APOE4 effects, we employed the KRAB-mediated CRISPRi gene regulation technique^67^ in young E4-KI mice to validate their functional significance by altering the expression of selected DEGs toward their expression levels in E3-KI mice. We selected two candidate genes based on our snRNA-seq results: Nell2 (neural epidermal growth factor-like protein 2), the only DEG shared between CA2/3 PC cluster 6 and DG cluster 2 (Supplementary Table 6), and Mdga2 (MAM domain containing glycosylphosphatidylinositol anchor 2), the only DEG shared between DG clusters 1 and 2 (Supplementary Table 6). We designed custom recombinant lentiviruses targeting each selected DEG for CRISPRi as well as dCAS9/KRAB vector lentivirus that would also induce mCherry expression for fluorescence tagging of target neurons, and injected them into the DG/CA3 hippocampal region in 5-month-old female E4-KI mice. Two months later, we recorded CA3 PC and DGC electrophysiological parameters in hippocampal slices from the injected mice. Using mCherry fluorescence as a guide, we were able to target both transduced CRISPR-positive and control CRISPR-negative neurons in the same slice for patch-clamp recordings (Fig. 8).

**Figure 8.**
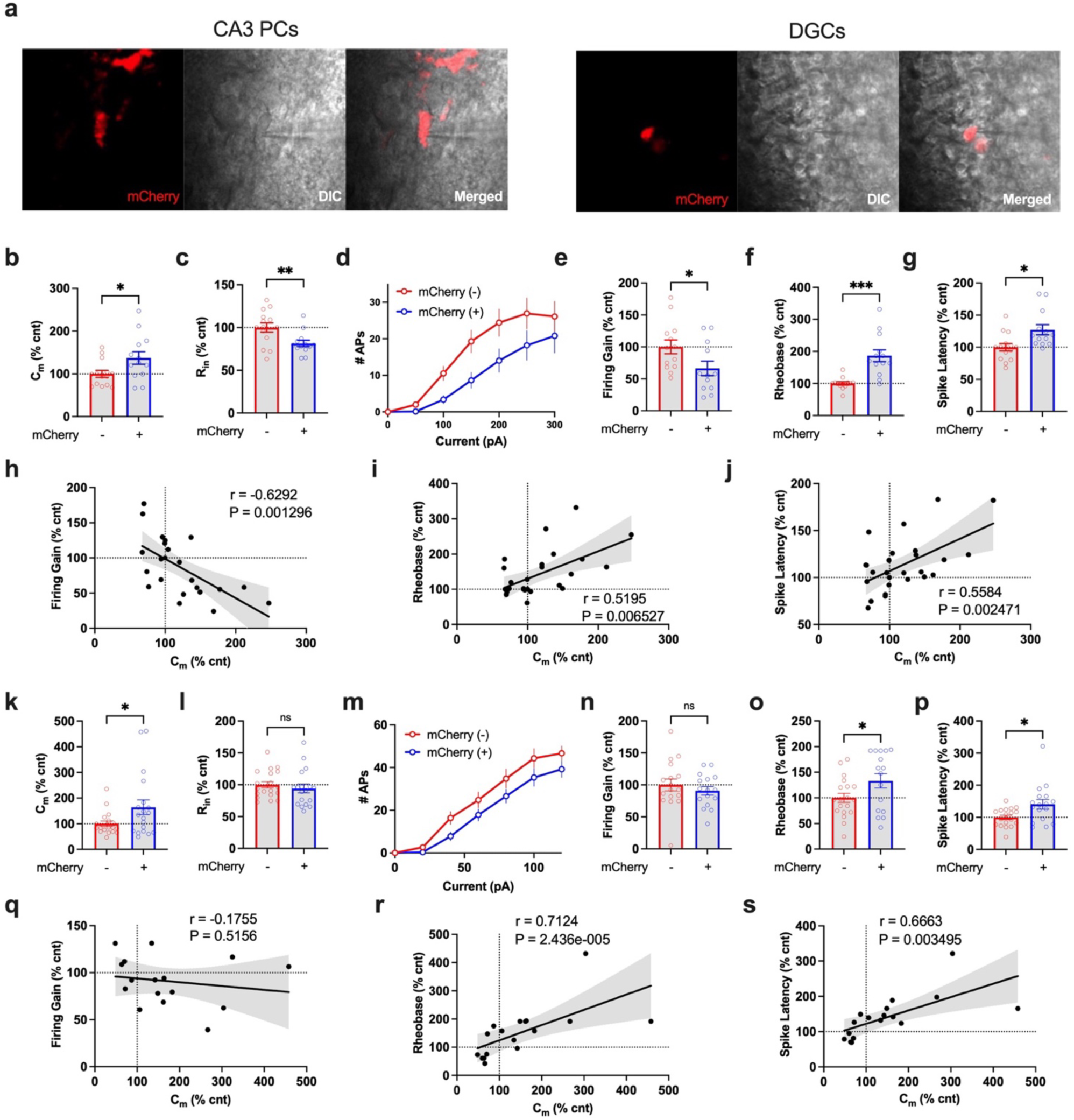
CRISPRi-mediated reduction in Nell2 expression normalizes cell soma size and excitability of transduced CA3 PCs and DGCs from young E4-KI mice. **a,** Example images of transduced cells expressing mCherry (red, left panels) targeted for whole-cell patch recording in acute brain slices under differential interference contrast (DIC) imaging (middle panels). Right panels: overlay. **b–g,** Nell2 CRISPRi/mCherry+ CA3 PCs display normalized morpho-electric and excitability parameters: C_m_ (b), R_in_ (c), I-F Curves (d), firing gain (e), rheobase (f), and spike latency (g). **h–j,** Increased C_m_ correlates with reduced excitability in all recorded CA3 PCs: C_m_ versus Firing gain (h), C_m_ versus Rheobase (i), and C_m_ versus spike latency (j). **k–p,** Nell2 CRISPRi/mCherry+ DGCs display normalized morpho-electric and excitability phenotype: C_m_ (k), R_in_ (l), not in I-F Curves (m) (see also Fig. 3b) or firing gain (n) (see also Fig. 3k,p), in rheobase (o), and spike latency (p). **q– r,** Increased C_m_ correlates with reduced excitability in all recorded DGCs, except in C_m_ versus Firing gain (q), C_m_ versus Rheobase (i), and C_m_ versus spike latency (j). All data are represented as mean ± s.e.m., *p<0.05, **p<0.01, ***p<0.001, unpaired two-tailed t-test or Mann-Whitney U test, or Pearson correlation analysis (two-sided). N= 5 and 4 for CA3 PCs and DGCs, respectively, from E4-KI mice.

Nell2 is a kinase C-binding glycoprotein^68,69^ that has been shown to have multiple roles in regulating neuronal proliferation, survival, differentiation, polarization, as well as axon guidance and synaptic functions^70–72^. mCherry+ Nell2-CRISPRi-transduced CA3 PCs and DGCs (Fig. 8a) had higher C_m_ values (Fig. 8b,k) and displayed significantly higher rheobase and spike latency values (Fig. 8f,g,o,p) compared to neighboring mCherry-non-transduced neurons, indicating reduced excitability of the Nell2-CRISPRi-transduced neurons. mCherry+ CA3 PCs also showed decreased firing gain (Fig. 8d,e), indicating a rescue of that cell type-specific phenotype, although we observed no differences in firing gain between mCherry+ and mCherry-DGCs (Fig. 8m,n), consistent with the lack of APOE4-induced gain phenotype in those neurons (Fig. 2c,d). Importantly, changes in C_m_ correlated significantly with changes in each of the three phenotypic IE parameters (Fig. 8h–j,r,s), supporting our conclusion that APOE4 reduced neuronal size and increased excitability phenotypes are related.

Mdga2 is a gene previously implicated in ASD-related epilepsy^73^. We found no difference in any morpho-electric and excitability parameters in mCherry+ Mdga2-CRISPRi-transduced neurons versus mCherry-DGCs (Extended Data Fig. 9). Although these data do not support involvement of Mdga2 in APOE4-induced neuronal morpho-electric phenotype, they do confirm that the CRISPRi procedure itself did not affect neuronal parameters and therefore can serve as a “negative” CRISPRi control. Altogether, our CRISPRi validation study confirmed the critical role of Nell2 overexpression in mediating early APOE4-induced neuronal morpho-electric deficits and substantiates the targeted reversal of APOE4-induced transcriptomic changes, such as reducing Nell2 expression, as a promising strategy for early therapeutic intervention in APOE4-related AD.

## Discussion

Multiple studies on young *APOE4* carriers have consistently demonstrated increased task-related activation of the hippocampus^25,74,75^. Notably, our data on early network hyperexcitability in cognitively normal young E4-KI mice being predictive of spatial cognitive impairment with aging is consistent with the reports that the degree of APOE4-induced hippocampal hyperactivity in young APOE4 carriers predicts future cognitive decline in humans^25^. Furthermore, we have previously found that reducing network hyperactivity in 9-month-old E4-KI mice via 6-week treatment with the GABA_A_ receptor potentiator pentobarbital prevented learning and memory deficits in these mice at 16 months^42^, supporting that early network hyperexcitability is a major trigger and promoter of APOE4-induced cognitive dysfunction with aging. Finally, our finding of region-specific network hyperexcitability in CA3 and DG is also consistent with reports showing greater hippocampal activation in young and non-demented aged APOE4 carriers^74,76^ as well as with studies on amnesic MCI patients showing hippocampal hyperactivation and atrophy specifically in CA3 and DG in humans^77^.

In APP-overexpressing AD mice, network hyperexcitability is underlain by the hyperexcitability of excitatory neurons^15,17,78^ as well as by inhibitory deficit^33,79^ resulting from Aβ toxicity. Multiple intrinsic excitability parameters mediating the Aβ effect on neuronal hyperexcitability have been reported in a number of transgenic amyloidosis mouse models, including depolarized neuronal resting membrane potential^15,17^, depolarized reversal potential of GABA-mediated inhibitory synaptic currents^17^, increased firing gain^80,81^, reduced spike latency^81^, as well as altered AP width^58,81^ and amplitude^58^. We found that CA3 PCs and DGCs in young E4-KI mice are hyperexcitable in their subthreshold (rheobase, spike latency) and suprathreshold (firing gain, max firing rate, firing accommodation) parameters. The observed APOE4-driven hyperexcitability phenotype resembles those reported in mouse amyloidosis models in some parameters but differs in others. For example, we found no changes in resting membrane potential or in the AP shape, indicating both overlapping and differential mechanisms underlying Aβ and APOE4-induced hippocampal hyperactivity in AD pathogenesis.

Neuronal excitability is also mediated by passive neuronal properties, such as cell size. In our study, CA3 PCs and DGCs in young E4-KI mouse hippocampus were consistently smaller than those in age-matched E3-KI mouse hippocampus, as evidenced by lower C_m_ values and confirmed by morphometric analysis of cell soma volumes. Our results are consistent with previous reports: APOE4 is associated with smaller neuron size in resected tissue from human epilepsy patients^82^. Neuronal APOE4 expression in mice results in reduced dendritic morphology^65^ and has been shown to exert a disruptive effect on dendritic integrity in response to excitotoxic injury^83^. Reduced C_m_ has also been reported in human induced pluripotent stem cell (iPSC)-derived neurons from AD patients^58^ as well as in neurons from APP/PS1 mice^59^. In CA3 PCs and DGCs from young E4-KI mice, every IE parameter that differed from those in E3-KI mice correlated significantly with C_m_, while no such correlation was observed in those neurons in E3-KI mice, suggesting that reduced cell size at least in part underlies APOE4-induced neuronal hyperexcitability. Previous studies on different AD mouse models reported a similar relationship: decreased neuronal size was shown to underlie hyperexcitability and hyperactivity of CA1 PCs in APP/PS1 mouse model of AD^59^; analogous causality has been reported in human iPSC-derived neurons carrying PS1 and APP mutations^58^.

In the brain, APOE is primarily expressed by astrocytes, but in response to stress or injury neurons also begin to express APOE^84–86^. Neuronal APOE4 can cause multiple pathologies including tau hyperphosphorylation, mitochondrial impairment, neuroinflammation, and neurodegeneration^63,85,87–92^. Neuronal APOE4 expression is also responsible for the impairment and death of hilar somatostatin-expressing interneurons, which correlates with learning and cognitive deficits in aged E4-KI mice^8,9,64^. However, the exact functional consequences of APOE4 expression in hippocampal excitatory neurons have not been demonstrated until now. Here we show that APOE4’s detrimental effects on neuronal size and excitability are specific to its expression in neurons: removal of APOE4 in neurons of young fE4-KI^Syn1-cre+^ mice normalized CA3 PC excitability as well as neuronal size and input resistance to levels similar to those in E3-KI mice, while astrocytic APOE4 removal in fE4-KI^GFAP-cre+^ mice showed no such rescue.

Our clustering analysis revealed that not all cells, even within a given neuron type, are equally vulnerable to APOE4’s detrimental effect on neuronal size and excitability. We uncovered two distinct subpopulations of neurons within each neuronal cell type: healthier, “normal” neurons as well as smaller and hyperexcitable ones, with the enrichment of each subpopulation dependent on neuronal cell type, APOE genotype, and age. This is consistent with our recent study^62^ showing that a subpopulation of neurons upregulates their APOE4 expression, contributing to selective cellular vulnerability to AD-related neurodegeneration. The same study showed that neuronal APOE expression levels are also age-dependent, with the high APOE4-expressing neuron population peaking at 10 months of age, while high APOE3-expressing neurons reached maximum at 15 months of age. This is comparable with what we observed in the current study, with many CA3 PCS and DGCs shifting into the hyperexcitable subpopulation earlier in young E4-KI and later in aged E3-KI mice, suggesting that heterogenous APOE4 expression in neurons might drive electrophysiological heterogeneity during aging, which warrants further in-depth study.

Curiously, CA3 PCs and Type II DGCs in aged E3-KI mice “catch up” with those from E4-KI mice, becoming smaller and more excitable, presumably as the result of aging^93–95^, thus abolishing the early morpho-electric difference in aged E4-KI and E3-KI mice. The fact that unlike E4-KI mice, aged E3-KI mice do not develop network hyperexcitability in those regions, even though their neurons become more excitable, suggests that additional mechanisms are involved in promoting the APOE4-induced network dysfunction at more advanced ages. We found that the population of smaller and hyperexcitable Type I DGCs was significantly more enriched in aged E4-KI mice, likely contributing to the maintained overall DGC hyperexcitability phenotype compared to E3-KI animals. Additionally, we also observed a progressive inhibitory deficit specifically in the DG of E4-KI mice with aging, characterized by a decreased frequency of spontaneous inhibitory inputs. This led to a significantly higher E-I ratio in aged E4-KI mice versus age-matched E3-KI mice, further exacerbating APOE4-induced network and behavioral dysfunction in advanced age^8,9,96,97^. This inhibitory deficit serves as a crucial link between early hippocampal hyperexcitability and impaired cognition in old age. Network hyperactivity places stress on APOE4-expressing interneurons through various mechanisms, including excitotoxicity and metabolic stress^34^. Neuronal APOE4 expression has been shown to result in neuronal loss following excitotoxic challenges^98^, and interneurons have been shown to be uniquely vulnerable to such insults owing to higher energy demand^99,100^ and lower resistance to chronic stress^101^. As noted above, our earlier study demonstrated this connection: arresting network hyperactivity with 6-week pentobarbital treatment in 9-month-old E4-KI mice prevented DG interneuron death and subsequent learning impairment observed 7 months later^42^, supporting that early APOE4-induced network hyperactivity likely drives interneuron depletion and contributes to cognitive impairment in aged E4-KI mice.

The exact mechanism behind the APOE4-induced neuronal atrophy and associated hyperexcitability remains unclear. However, despite the radically different pathological triggers, our observations of similar neuronal atrophy and hyperexcitability in E4-KI mice lacking Aβ accumulation and of those reported in mouse amyloidosis models, imply a shared pathway of action. One such pathway may involve tau^102,103^: neuronal APOE4 can induce an increase in tau phosphorylation^104^ and neurodegeneration^65^, mirroring the Aβ effects on tau reported in other studies^105–108^. Moreover, removal of endogenous tau prevented neuronal loss and cognitive deficits in both E4-KI mice^8^ and APP transgenic mice^109,110^, and we recently reported that removal of neuronal APOE4 is protective against tau pathology, network hyperexcitability, neurodegeneration, and neuroinflammation in a tauopathy mouse model^63^.

Another of our recent studies highlighted the potential therapeutic benefits of reversing APOE4-induced transcriptomic changes on ameliorating network and cognitive deficits in APOE4 models of AD^111^. Here, our snRNA-seq analysis revealed a number of genes that exhibited differential expression patterns that mirrored the observed APOE4-specific and aging-dependent electrophysiological changes; these DEGs may influence neuronal morphology and excitability directly, or alternatively regulate the susceptibility or resilience to neuronal APOE4 toxicity. Importantly, we identified a specific overexpression of Nell2 in CA3 PCs and DGCs of young E4-KI mice, which was abolished upon neuronal removal of APOE4. CRISPR_i_ validation study provided proof for the critical role of Nell2 overexpression in mediating APOE4-induced early neuronal morpho-electric deficits, as reducing Nell2 expression in neurons rescued neuronal size and hyperexcitability phenotype in both CA3 PCs and DGCs of young E4-KI mice.

Nell2 is a glycoprotein that is specific to glutamatergic neurons in pyramidal and granule cell layers in CA1–3 and DG, respectively^72^. It has been shown to interact with protein kinase C through its EGF-like domain^68^ and to modulate MAPK activity^70^. Furthermore, Nell2 shares domains with thrombospondin-1, a protein involved in regulating cell proliferation and apoptosis^112^. It is secreted as a trophic factor^70,71^, however its alternative splicing variant is cytoplasmic and was shown to be involved in intracellular signaling^113^ (such as binding to PKCβ1^68^). Our results suggest a cell-autonomous (and therefore cytoplasmic variant) role of Nell2 signaling in mediating APOE4 effects, as Nell2 reduction in CRISPRi-transduced neurons lowered their excitability, while neighboring, non-transduced neurons remained hyperexcitable. Nell2 is one of 65 genes in the “neuron cellular homeostasis” Gene Ontology pathway (GO:0070050), alongside known AD-associated genes such as APP, PSEN1 and PSEN2, TMEM106B^114^, SLC24A2^115^ and IL-6^116^. Increased NELL2 protein expression was observed in the prefrontal cortex of AD patients^117^, in AD patient iPSC-derived hippocampal neurons^118^, and in the hippocampus of 5XFAD AD model mice^119^. A recent study analyzing CSF biomarkers in cognitively unimpaired individuals over the age of 65 identified Nell2 to be one of five proteins whose CSF levels significantly associated with both total tau and p-tau levels^120^. Additionally, NELL2 circular RNA (circRNA) expression was found to be upregulated in cortical tissue of AD patients^121^. Clearly, the roles of Nell2 in neuronal morphology and/or excitability, especially in the context of APOE4, warrants further in-depth study.

In summary, our study provides evidence that the expression of APOE4 in excitatory hippocampal neurons of young E4-KI mice induces significant gene expression changes, including overexpression of Nell2 in CA3 PCs and DGCs, leading to neuronal atrophy and hyperexcitability in specific neuronal subpopulations and resulting in early network hyperexcitability that predicts future cognitive impairment. These findings suggest a causative timeline of APOE4-induced network dysfunction with aging: during early stages, neuronal APOE4 expression triggers cell type-specific transcriptomic alterations, neuronal atrophy, and hyperexcitability, culminating in network hyperactivity. As the E4-KI mice age, this process further contributes to the loss of GABAergic interneurons and inhibitory deficits, accompanied by a pathological increase in hyperexcitable Type I DGCs, leading to network failure and cognitive impairment. In contrast, aged E3-KI mice display resistance to cognitive decline due to maintained inhibitory function despite increased excitability of primary neurons. To develop effective intervention strategies for APOE4-related AD, further research is needed to unravel the precise mechanisms underlying APOE4-induced, cell type- and age-dependent changes in excitatory and inhibitory neurons in the hippocampus. Deepening our understanding of these processes holds the potential to pave the way for targeted therapeutic interventions for APOE4-related AD.

## Methods

### Mice

Human E3-KI and E4-KI mice^122^ were from Taconic. LoxP-floxed APOE knock-in (fE-KI) mice were generated in our lab as described previously^123^. Mice with a conditional deletion of the human *APOE* gene were created as described previously^64^: homozygous fE3-KI and fE4-KI mice were crossbred with GFAP-Cre transgenic mice [B6.Cg-Tg(GFAP-cre)8Gtm]^124,125^ or Synapsin 1-Cre (Syn1-Cre) transgenic mice [B6.Cg-Tg(Syn1-cre)671Jxm/J]^126^. These crosses produced mice that were heterozygous for APOE3 or APOE4 and positive for GFAP-Cre or Syn-1-Cre. These mice were further crossbred with fE3-KI or fE4-KI mice to generate mice that were homozygous for floxed APOE3 or APOE4 and positive for GFAP-Cre (fE3-KI^GFAP-Cre+^ or fE4-KI^GFAP-Cre+^) or Syn1-Cre (fE3-KI^Syn1-Cre+^ or fE4-KI^Syn1-Cre+^). Littermates negative for GFAP-Cre or Syn1-Cre were used as controls. All mice were on a pure C57BL/6 genetic background and were housed in a pathogen-free barrier facility on a 12 h light cycle at 19-23°C and 30-70% humidity. Animals were identified by ear punch under brief isoflurane anesthesia and genotyped by polymerase chain reaction (PCR) of a tail clipping. All animals otherwise received no procedures except those reported in this study. All animal experiments were conducted in accordance with the guidelines and regulation of the National Institutes of Health, the University of California, and the Gladstone Institutes under the protocol AN176773.

### Viral injections

For viral transduction, 5-month-old female mice were anesthetized with an initial intraperitoneal (IP) injection of ketamine/xylazine mix and maintained with 1% isoflurane oxygen mix at a flow rate of one liter per minute. The scalp area was cleaned and hair chemically removed. A small incision in the scalp was made to reveal the midline suture from bregma to lambda. Using a stereotactic frame equipped with a small high-speed drill 0.5mm burr holes were drilled through the skull at coordinates 2.1mm posterior to bregma and ±1.5mm lateral to the midline, at a depth to reveal the surface of the cortex. A 28 gauge canula attached to a syringe pump was lowered to a depth of 1.5mm below the cortical surface and 2uL of lentivirus was injected at a rate of 500nL/min. After each delivery we waited an additional 2 minutes for pressure equalization before the canula was slowly removed. The scalp was sutured closed with silk thread. At the end of surgery mice were treated with subcutaneous ketofen injection near the incision site and IP injections of buprenorphine and one of saline for hydration. Mice were allowed to recover in new cages warmed by a heating pad. At the end of the four-hour recovery a second injection of buprenorphine was administered and cages were returned to primary housing racks.

### Tissue slice preparation

Mice were deeply anesthetized with isoflurane and decapitated. The brain was rapidly removed from the skull and placed in the 4°C slicing solution comprised of 110 mM Choline Chloride, 2.5 mM KCl, 1.25 mM NaH_2_PO_4_, 26 mM NaHCO_3_, 2 mM CaCl_2_, 1.3 mM Na Pyruvate, 1 mM L-Ascorbic Acid, and 10 mM dextrose. 300μm sagittal sections were cut using a vibratome (VT 1200s, Leica). Following slicing, slices were transferred into a vapor interface holding chamber (Scientific Systems Design Inc, Canada) aerated with 95% O_2_/5% CO_2_ gas mixture and allowed to recover at 34°C for 1 h before recording.

### Immunohistochemistry

APOE-KI mice were analyzed at 5 and 10 months of age. Brain tissue collection occurred after the mice were administered intraperitoneal injections of avertin (Henry Schein) and transcardially perfused with 0.9% saline for 1 minute. Right hemispheres were drop-fixed for 48 hours in 4% paraformaldehyde (Electron Microscopy Sciences) and then placed in 1× PBS (Corning). Brains were then switched to 30% sucrose for 48 hours before mounting onto a freeze sliding microtome (Leica) and sectioning into 30um slices. Sections were stored in cryprotectant solutions at −20°C (30% ethylene glycol, 30% glycerol, and 40% 1× PBS). Sections selected for staining were then rinsed thrice with x 1× PBS-T (PBS with 0.1% Tween-20) (Millipore Sigma) and incubated for 10 minutes in boiling antigen retrieval buffer (Tris buffer, pH 7.6; TEKNOVA). Next, sections were again rinsed in 1× PBS-T before being incubated for an hour in a blocking solution (5% normal donkey serum (Jackson Labs), 0.2% Triton-X (Millipore Sigma) in 1× PBS), followed by an hour in Mouse-on-Mouse (MOM) Blocking Buffer (one drop MOM IgG in 4 ml PBS-T) (Vector Labs). Following the MOM block, the sections were incubated overnight at 4 °C with the primary antibody, which was diluted to an optimal concentration of 1:500 for anti-NeuN (Millipore Sigma). The next day, sections were rinsed thrice with 1× PBS-T and then incubated for 1 hour in secondary antibody (donkey anti-guinea pig 594, 1:1000; Jackson Immuno) with DAPI (1:20,000). After this, the sections were washed again in PBS-T, mounted on microscope slides (Fisher Scientific), and coverslipped using ProLong Gold mounting medium (Vector Laboratories). Images were captured using an FV3000 confocal laser scanning microscope (Olympus) at ×40 magnification. To minimize batch-to-batch variation, all samples were stained simultaneously and imaged at the same fluorescent intensity.

### Cell soma volume analysis

To measure cell soma volumes from stained sections, we used the python implementation of the Cellpose software^127^. We applied this tool to identify candidate neuron and nuclear masks in the CA3 cell layer. To ensure high-quality soma reconstructions, we employed four filtering criteria: 1) Boundary Exclusion: Cell masks located on the edges of the field of view were removed to eliminate partial cells; 2) Z-Plane Consistency: Neurons detected in fewer than two z-planes were excluded to avoid partially captured cells within the field of view; 3) Nuclear Integrity: Masks containing more than one nucleus were discarded, as they likely represented merged cells; and 4) Shape Regularity: We required that the ratio of the mask volume to the volume of its bounding box exceed 0.4, retaining only masks with regular shapes and eliminating significant non-neuronal background signals. Applying these criteria, we obtained 141 and 114 well-defined neurons from 6 young E3-KI and 6 young E4-KI mice, respectively.

### Whole-cell patch clamp electrophysiology

Brain slices were placed into a submerged dual-side recording chamber (RC-27D, Warner Scientific) and perfused at 5 mL/min with oxygenated ACSF solution at 34°C. Recording and holding ACSF solution was comprised of 124mM NaCl, 26mM NaHCO_3_, 10mM Glucose, 1.25mM NaH_2_PO_4_, 2.5mM KCl, 1.25mM MgCl_2_, and 1.5mM CaCl_2_. Neurons were imaged using a modified Olympus BXW-51 microscope with a 60x objective (Scientifica Inc, Great Britain). Whole-cell patch-clamp recordings were performed using a Multiclamp 700B amplifier with signals sampled at 10 kHz and digitized using a Digidata 1550B with Axon pCLAMP software (all from Molecular Devices, USA).

#### Intrinsic neuronal excitability and passive parameter measurements

Cells were allowed to stabilize for minimum of 3 minutes after obtaining whole-cell patch configuration before recordings commenced. Patch pipettes were filled with potassium-gluconate based solution containing 122.5 mM K-gluconate, 8 mM KCl, 10 mM HEPES, 2 mM MgCl_2_, 0.2 mM EGTA, 2 mM ATPNa, and 0.3 mM GTPNa, pH set to 7.2–7.3 with KOH and with osmolarity of 270-280 mOsm. To block spontaneous synaptic activity, AMPA receptor antagonist 6-cyano-7-nitroquinoxaline-2,3-dione (CNQX, 20μM), NMDA receptor antagonist D--2-amino-5-phosphonopentanoic acid (AP-5, 50μM) and GABA_A_ receptor antagonist Picrotoxin (5μM) were added to the perfusate.

#### Cell membrane capacitance (C_m_)

C_m_ was measured in voltage clamp configuration at -70mV using Clampex software’s Membrane Test protocol as the integral of the fast capacitance transient current resulting from a 10mV, 5ms square pulse stimulation.

#### Input resistance (R_in_)

R_in_ was measured in current clamp configuration at V_m_ of -70mV using 1s, 10pA current injections from -50 to +50pA. Resulting I-ι1V_m_ curve was fitted with a linear slope to calculate the input resistance. *Resting Membrane Potential* was measured in current clamp configuration at I=0.

#### Input current-spiking frequency (I-F)

I-F relationship was recorded in current clamp configuration using incremental 1-s 20pA (CA1 PCs and DGCs) or 50pA (CA3 PCs) depolarizing current injections at 0.1Hz with resting membrane potential held at -70mV. Resulting spike count was then plotted against the current step magnitude to generate the I-F curve.

#### Firing gain

Firing gain was calculated as the linear slope of the first 5 steps in the I-F curve starting from rheobase.

#### Rheobase

Rheobase was determined from -70mV as the minimum magnitude of the 1s depolarizing current injection required to elicit an action potential.

#### Spike latency

Spike latency was measured in current clamp configuration at -70mV using a 1s 800pA current ramped injection, as the delay from the beginning of the stimulation to the generation of the first action potential (when dV_m_ reached 20V/s).

#### Accommodation

Accommodation was measured as the ratio of last to first interspike interval in response to a 1s step current injection at twice the rheobase.

#### Action potential threshold

Action potential threshold was measured at rheobase step as the V_m_ value where dV_m_ crossed 20V/s.

#### Fast afterhypolarization (fAHP)

fAHP was measured as a negative-going peak following the action potential relative to the action potential threshold under a 1s current step injection that elicited at least 4 action potentials.

#### Slow afterhypolarization (sAHP)

sAHP was measured as the negative-going peak relative to baseline within a 1s period immediately after 1s 300pA current step injection.

#### Spontaneous synaptic activity recording

Postsynaptic currents were recorded using a Cs^+^-based intracellular pipette solution containing: 140mM CsMeSO_4_, 10mM HEPES, 2mM MgCl_2_, 0.6mM EGTA, 2mM ATPNa, 0.3mM GTPNa, set to pH 7.2–7.3 with CsOH and with osmolarity of 270-280 mOsm. Spontaneous excitatory postsynaptic currents (sEPSCs) were recorded at -70mV, inhibitory postsynaptic currents (sIPCSs) at 0mV, for a period of 3-5 minutes.

### Single-nuclei preparation for 10x loading

For analysis of transcriptome in various types of cells in the hippocampus from E3-KI and E4-KI mice at different ages, snRNA-seq dataset from our previous publication^62^ (GEO accession: GSE167497) was used for the analysis in the current study.

### Single-nuclei preparation from 5- and 10-month-old fE4-KI^Syn1-cre+^ mice

To isolate single nuclei from adult mouse brains, we combined and adapted the 10x Genomics-demonstrated protocol for nuclei isolation from adult mouse brain^128^ and the Allen Brain Institute protocol for fluorescence-activated cell sorting (FACS) of single nuclei56 as follows. Hippocampi were acutely dissected on ice. Dissected hippocampi were placed in 2-ml Hibernate A/B27/GlutaMAX (HEB) medium in a 5-ml tube. The HEB medium was removed to a 15-ml conical tube and kept on ice. Next, 2 ml of chilled lysis buffer (10 mM Tris-HCl, 10 mM NaCl, 3 mM MgCl2 and 0.1% Nonidet P40 Substitute in nuclease-free water) was added to the tissue, and the hippocampi were homogenized by suctioning ten times through a 21-gauge needle. After homogenization, the tissue was lysed on ice for 15 min, swirling 2–3 times during this incubation period. The reserved chilled HEB media was then returned to the lysed tissue solution, and the tissue was further triturated with 5–7 passes through a 1-ml pipette. A 30-µm cell strainer (MACS SmartStrainer, Miltenyi Biotec, 130110-915) was washed with 1 ml of PBS, and the lysed tissue solution was filtered through the strainer to remove debris and clumps. Filtered nuclei were centrifuged at 500 relative centrifugal force (r.c.f.) for 5 min at 4 °C. The supernatant was removed, and nuclei were washed in 1 ml of nuclei wash and resuspension buffer (1× PBS with 1.0% BSA and 0.2 U µl−1 of RNase inhibitor). Nuclei were again centrifuged at 500 r.c.f. for 5 min at 4 °C and resuspended in 400 µl of nuclei wash and resuspension buffer. DAPI was added to a final concentration of 0.1 µg ml−1, and the nuclei were filtered through a 35-µm cell strainer. DAPI-positive nuclei were sorted by gating on DAPI-positive events, excluding debris and doublets, using the BD FACSAria II at the Gladstone Institute’s Flow Cytometry Core.

### Complementary DNA library preparation and sequencing

cDNA libraries were prepared using the 10x Chromium Single Cell 3’ GEM, Library & Gel Bead Kit according to the manufacturer’s instructions. 10x Chromium v2 kit was used for the APOE-KI dataset. 10x Chromium v3 kit was used for APOE-fKI/Syn1-Cre dataset (10x Genomics: 1000092). Libraries were sequenced on an Illumina NovaSeq 6000 sequencer at the UCSF Center for Advanced Technology Core.

### Pre-processing and clustering of 5- and 10-month-old fEKI^Syn1-cre+^ mouse snRNA-seq samples

Clustering was performed following a previously established method^62^. Demultiplexed FASTQ files were aligned to a custom reference genome built from mm10-1.2.0, which includes introns. This alignment was conducted using the Cell Ranger v2.0.1 counts function with default parameters, as outlined in the Cell Ranger documentation. UMI counts were also determined using the Cell Ranger counts function, and count matrices from multiple samples were merged into a single count matrix using Cell Ranger’s aggr function with default parameters. Barcodes (potential cells) were filtered based on a UMI count threshold. The filtered UMI count matrices were further processed using Seurat v2.3.4. Specifically, the data were filtered to include only protein-coding genes. Cells were filtered to include only those with 200-2,400 detected genes, 500-4,500 UMIs, and less than 0.25% mitochondrial reads, ensuring data quality. To normalize the gene expression matrices, the Seurat NormalizeData function^129,130^ was employed with a scale factor of 10,000. Clustering was performed using the implementation in Seurat v2.3.4. This algorithm involved embedding cells in a k-nearest neighbor graph based on Euclidean distance in PCA space. The edge weights between cells were refined using Jaccard similarity. The Louvain algorithm was used for clustering. Highly dispersed genes were identified using the Seurat FindVariableGenes function^129,130^. These genes were selected based on an average expression range of 0.25-4 and a minimum dispersion of 0.55. Nearest neighbor distances were computed using up to the first 15 principal components and a resolution of 0.6, leading to the identification of 28 distinct clusters.

### Cell type assignment

Cell type assignment was done as described previously^62^: The t-SNE data visualization technique was used to examine the presence of clusters in which mouse ages and genotypes were intermingled, without any observable indications of batch effects associated with genotype or age. To identify marker genes for each cluster, the FindAllMarkers function in Seurat^129,130^ was utilized. This algorithm uses the Wilcoxon rank-sum test to iteratively compare gene expression within a putative cluster against the expression in all other clusters. Marker genes were selectively chosen to exhibit positive expression (i.e., higher expression in the cluster of interest compared to other clusters), to be detected in at least 10% of cells within the cluster, and to demonstrate a 0.25 log2 fold higher expression in the cluster of interest relative to other clusters. To determine broad cell classes, including excitatory and inhibitory neurons, astrocytes, oligodendrocytes, and OPCs, marker genes were compared against cell-type-specific markers obtained from previous RNA sequencing data on sorted cell types^131^. Moreover, for further subdivision of hippocampal cell types, particularly in the identification of principal cell subsets, marker genes for each cluster were compared against hippocampal cell-type-specific marker genes reported in hipposeq^132^. In the case of less common non-neuronal cell types, we compared gene expression in our cells with those genes enriched in each hippocampal cell type relative to all other cells in the hippocampus, as described in the DropViz resource^133^. To further corroborate the identity of the clusters, we additionally queried the top marker genes for each cluster against the Allen Brain’s genome-wide atlas of gene expression in the adult mouse brain^134^.

### Differential Gene Expression

To find genes that matched the selection criteria of interest based on the significant effects in our morpho-electric data, a set of differential gene expression tests were done between cell populations of interest. Statistically significant DEGs were detected using the non-parameteric Wilcoxon rank sum test, which was implemented using the FindMarkers function in Seurat. Only genes that are detected in a minimum fraction of 0.1 in either cell population were considered. A differential expression level threshold of absolute logFC = 0.05 was used. The p-value was adjusted based on Bonferroni correction. Genes with adjusted p-value < 0.05 were considered significantly differentially expressed. For the combined analysis of multiple differential expression comparisons, genes matching the differential expression criteria across multiple comparisons were chosen for further analysis.

### CRISPRi vector design

dCas9/KRAB-mediated CRISPRi vectors for Nell2 and Mdga2 were custom-designed, produced and packaged into lentivirus by VectorBuilder (USA) using previously published methods^67,135,136^. Specifically, multiple CRISPRi sgRNA sequences were designed for each target gene using VectorBuilder’s algorithm. For each target, two sgRNAs were chosen based on their off-target potential, proximity to the transcript start area, and sequence characteristics (excluding sequences with consecutive Cs and Gs). They were constructed into a lentivirus vector for the CRISPRi application.

### Single cell electrophysiology data analysis and statistics

Electrophysiological data analysis was done in Igor Pro 8 (Wavemetrics Inc, USA) using custom scripts. Postsynaptic events were quantified using a template matching algorithm in NeuroMatic v.3c plug-in for Igor Pro (Jason Rothman).

### K-means clustering analysis

Cluster analysis was performed using python (v3.10.12) scripts. For each cell type, clustering was performed with ages and genotypes combined. Electrophysiological measures were normalized in to z-scores. The K-means clustering algorithm^137^ implemented by the Scikit-learn package^138^ was applied to the complete set of normalized data as well as sub sets using five-fold cross validation where we randomly selected 80% of the neurons to use for cluster fitting, with the remaining 20% of the neurons classified based on the resulting clusters. This was repeated for 1000 iterations and the cluster centroids along with neuronal cluster assignment were recorded for each iteration. The probability of a cell maintaining its membership in the original cluster (Fig. 5c, lower panel) and the probability of each iteration to match the clustering of the full dataset was used to validate consistency of the clustering paradigm and to determine appropriate cluster number (k-value). To calculate strength of membership residuals, the distances from each centroid determined by the k-means algorithm were normalized to z-scores, and for each cell the difference (residual) in distances were used to estimate the affinity for Hyperexcitable or Normal clusters. These residuals for different genotypes and ages were compared with two-way ANOVAs and post-hoc Tukey tests.

### In vivo electrophysiology

For in vivo network excitability analysis, we re-analyzed raw LFP recording data from our previous study^11^. Briefly, data was collected from Neuronexus probes (configuration A4×8-400-200-704-CM32) which had four 5 mm shanks spaced 400 mm apart with 8 electrode sites per shank and 200 mm spacing between sites. Data from all mice were collected, amplified, multiplexed, processed, and digitized using a 32-channel Upright Headstage, commutator, and Main Control Unit (SpikeGadgets). Simultaneous data acquisition at 30 kHz and video tracking at 30 frames per second were conducted with Trodes software (SpikeGadgets). Each data collection period consisted of five days of 60-minute home cage sessions. To control for the effects of circadian rhythm, each mouse was recorded at a randomly assigned time each day across the light cycle. The anatomical location of each electrode site was determined by examining Nissl-stained histological sections and raw LFP traces. Data were down-sampled to 5kHz, high-pass filtered at 0.1Hz, and Gaussian smoothed at 2ms. To remove movement artefacts, all recordings were secondary-referenced to a channel located in corpus collosum^139^. LFP recordings were then analyzed using custom software written in MATLAB (Mathworks), incorporating the Chronux (http://chronux.org), Trodes to MATLAB (SpikeGadgets), and Neuroquery libraries, as well as Igor Pro 8 (Wavemetrics Inc, USA) with custom macros. Interictal spikes were detected in cell layers during periods of mouse activity as any event exceeding 5 SD above baseline for 10-100ms.

### Behavioral test

For learning performance evaluation, we re-analyzed Morris water maze (MWM) data from a longitudinal cohort of aged E3-KI and E4-KI mice in our previous study^11^. Briefly, mice were kept in the testing room with the arena hidden by a partition and given 2 days to acclimate to the environment in a new cage before starting the training. In each trial, mice were placed in a 122 cm diameter pool filled with opaque water and had to find a 14 × 14 cm platform submerged 1.5 cm below the water’s surface. The mice could only rely on spatial cues on the walls around the pool to guide their search. During the initial 2 days of pretraining, mice swam down a rectangular channel to locate the platform or were guided there by the experimenter after 90 seconds. After these sessions, the rectangular guides were removed, and the platform was moved to a new location. Over the next 5 days, mice underwent 4 daily hidden platform trials, where they were released from random locations and given 60 seconds to find the platform. These daily trials were divided into two pairs 10 minutes apart, with a 4-hour break between the pairs.

### Statistical analysis

Statistical analysis was performed using Prism 9 software (GraphPad Software LLC). To examine the significant effects of age and APOE genotype on neuronal excitability, we compared a maximum likelihood model that included age and APOE genotypes with a null model that disregarded them. The resulting p-values for each of the excitability parameters were adjusted for multiple comparisons using the false discovery rate method (FDR=5%). For parameters exhibiting significant age or APOE genotype effects, we performed focused comparisons between E3-KI and E4-KI mice, or between two different ages within each APOE genotype, utilizing either a student’s t-test or non-parametric tests, without applying any additional correction methods. Maximum Likelihood Test and false discovery rate correction were performed using custom scripts in python using the Scikit-learn^138^ and statsmodels^140^ packages. Exact statistical tests used and number of mice (N) are specified in figure legends.

## Data availability

All data associated with this study and the information of used materials are available in the main text, the Materials and Methods, or the Supplementary Information section. The snRNA-seq datasets of E3-KI and E4-KI mice at different ages are used from our previous publication^62^ (GEO accession: GSE167497). The snRNA-seq datasets of fE3-KI^Syn1-cre+^ mice that were generated during the study will be made available at GEO (accession # xxxxxxxx) upon acceptance of the paper. Data associated with all figures are also available as Source Data.

## Code availability

All code generated during this study is accessible via reasonable request to the corresponding authors’ lab.

## Acknowledgments

We thank Huang lab staff for their valuable discussions about the experimental design as well as data analyses and interpretation. We thank Reuben Thomas for his advice on statistical analysis methods. We also thank Eric Chow and the staff at the UCSF Center for Advanced Technology Core for advice and support with RNA sequencing, and Kathryn Clairborn for editorial assistance.

## Author Contributions

DT, SSJ, and MZ carried out most of the study and analyzed the data. EAAJ performed in vivo LFP recordings. BG, LD, and MZ performed snRNA-seq data analysis. OY, JB, NK, and ARZ participated in some experiments. Y. Hao did snRNA-seq of the hippocampi from fE4-KI^Syn1-Cre+^ mice. SYY conducted the MWM tests. SYY and SDL managed mouse lines. QX helped design CRISPRi vectors. MZ and YH designed and supervised the study. DT, MZ, and YH wrote the manuscript.

---

**Extended Data Figure 1.**
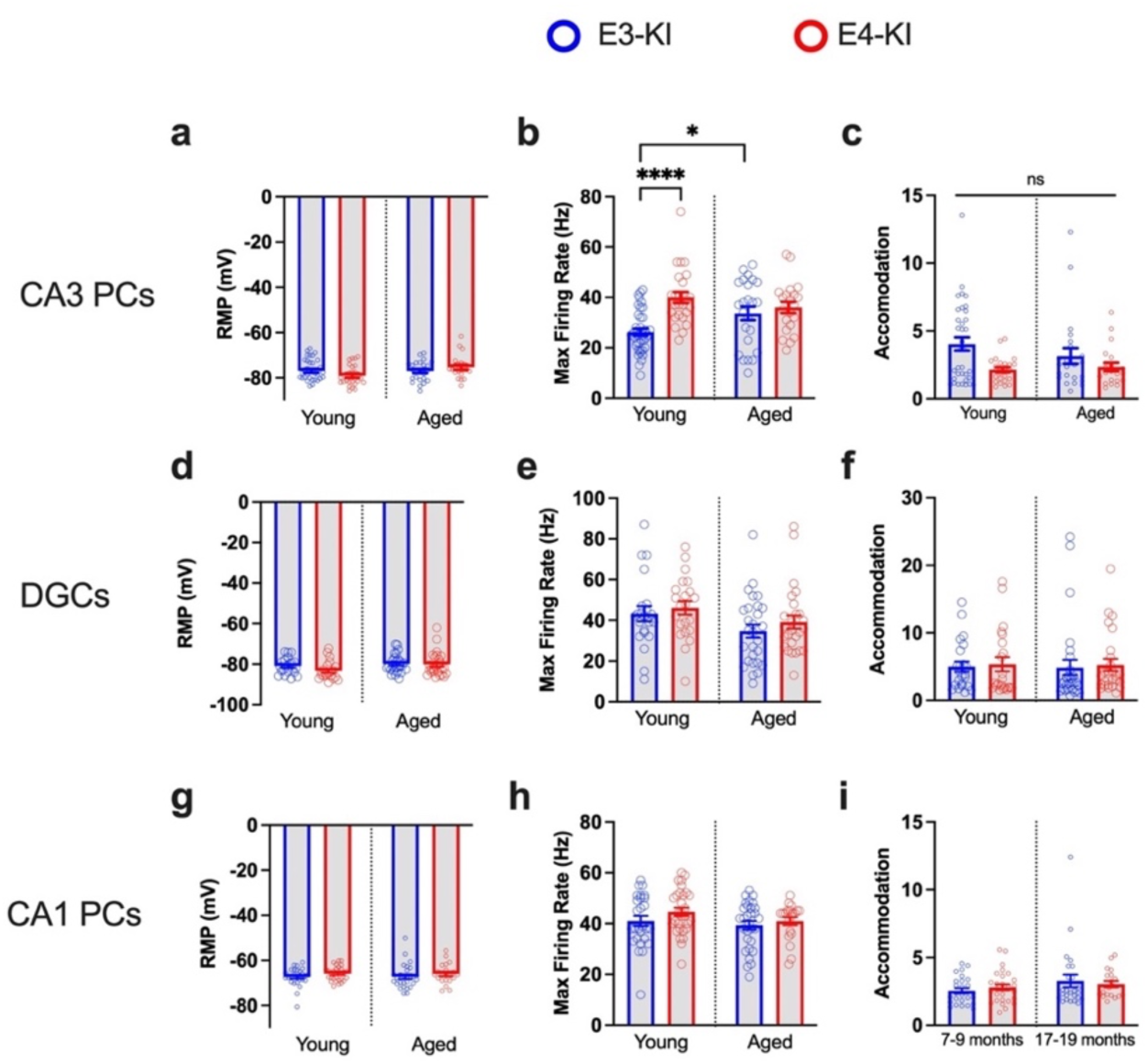
Secondary intrinsic excitability parameters in hippocampal excitatory neurons from E3-KI and E4-KI mice. **a–c,** CA3 PC IE parameter values: membrane potential (RMP) (a), maximal firing rate (b), and firing accommodation (c). **d–f,** Type II DGC IE parameter values: membrane potential (RMP) (d), maximal firing rate (e), and firing accommodation (f). **g–i,** CA1 PC IE parameter values: membrane potential (RMP) (g), maximal firing rate (h), and firing accommodation (i). All data are represented as mean ± s.e.m., *p<0.05, ****p<0.0001, unpaired two-tailed t-test or Mann-Whitney test.

**Extended Data Figure 2.**
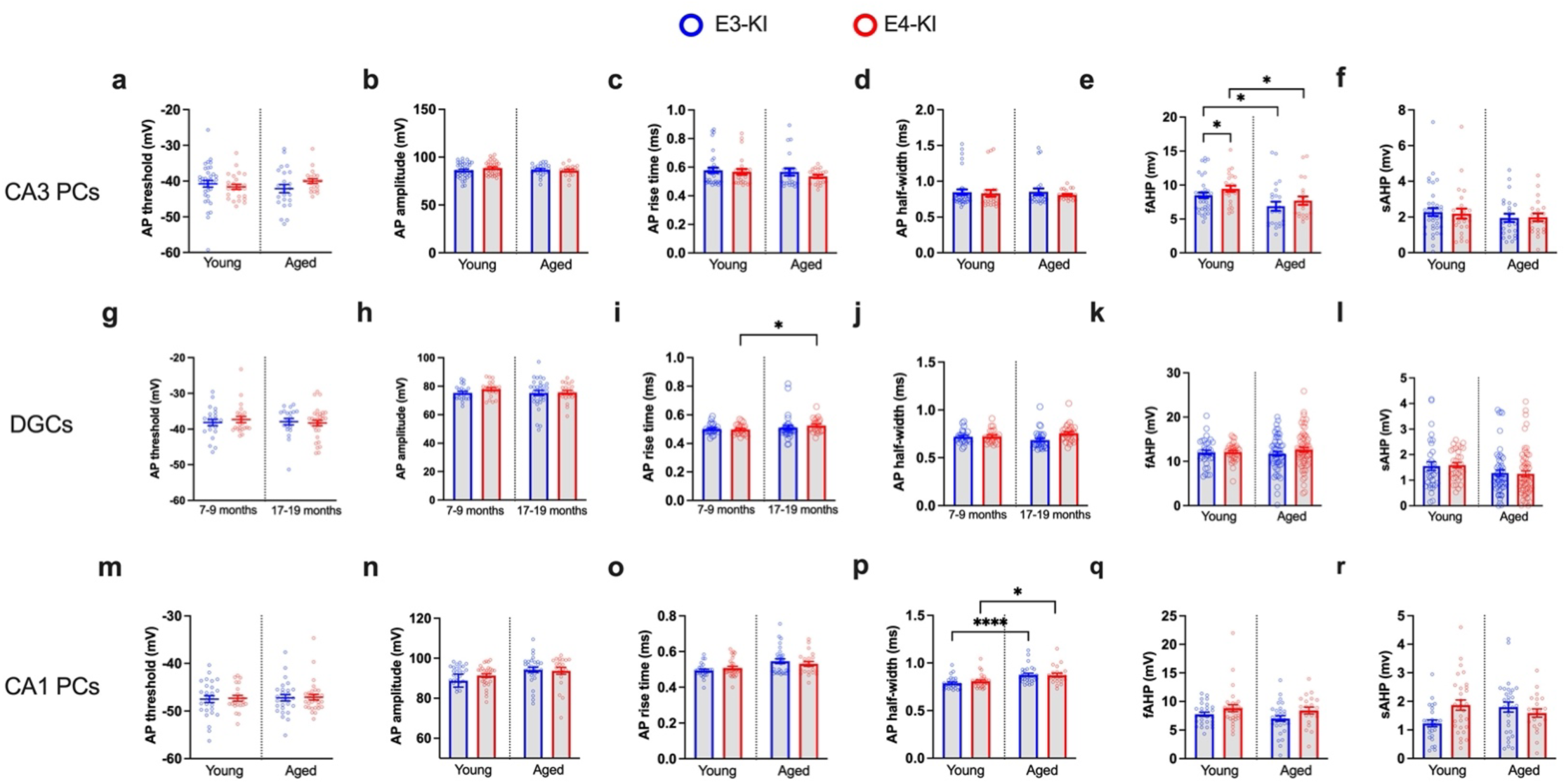
Action potential-related excitability parameters in hippocampal excitatory neurons from E3-KI and E4-KI mice. **a–f,** CA3 PC action potential (AP) waveform parameters: threshold potential (a), amplitude (b), rise time (c), half-width (f), fast afterhyperpolarization, fAHP (e), and slow afterhyperpolarization, sAHP (f). **g–l,** Type II DGC AP parameters: threshold potential (g), amplitude (h), rise time (i), half-width (j), fast afterhyperpolarization, fAHP (k), and slow afterhyperpolarization, sAHP (l). **m–r,** CA1 PC AP parameters: threshold potential (m), amplitude (n), rise time (o), half-width (p), fast afterhyperpolarization, fAHP (q), and slow afterhyperpolarization, sAHP (r). All data are represented as mean ± s.e.m., *p<0.05, ****p<0.0001, unpaired two-tailed t-test or Mann-Whitney test.

**Extended Data Figure 3.**
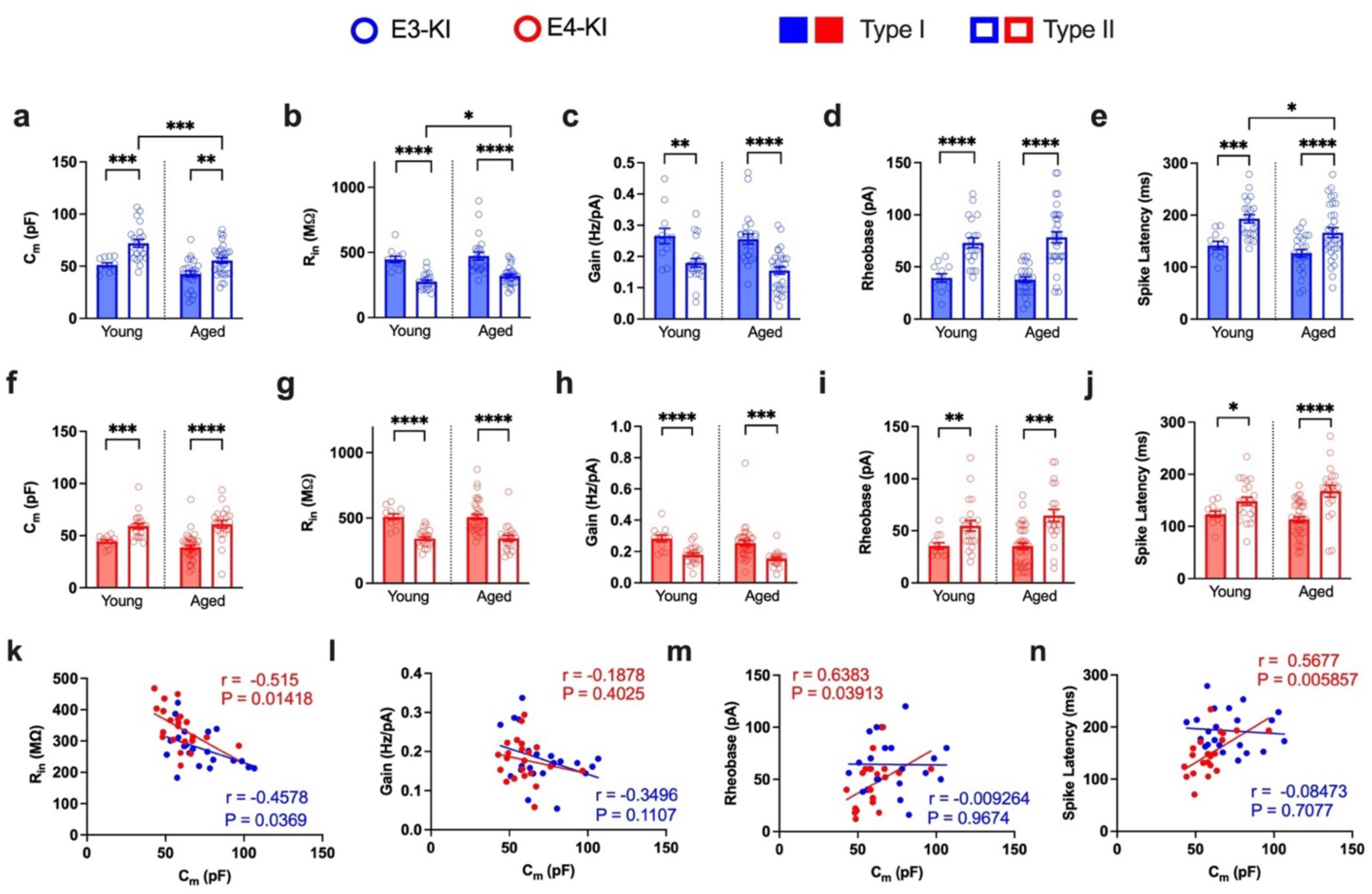
Morpho-electric characteristics defining the two dentate granule cell (DGC) subtypes in E3-KI and E4-KI mice. **a–e,** DGC characteristics for Type I (shaded bars) and Type II (clear bars) DGCs in E3-KI mice: C_m_ (a), R_in_ (b), firing gain (c), rheobase (d), and spike latency (e). **f–j,** DGC characteristics for Type I and Type II DGCs in E4-KI mice: C_m_ (f), R_in_ (g), firing gain (h), rheobase (i), and spike latency (j). **k,** C_m_ and R_in_ values correlate in Type II DGC from young E3-KI and E4-KI mice. **i,** Cm does not correlate with firing gain in Type II DGCs (see also Figure 3k) ; Pearson’s correlation analysis (two-sided). **m,** C_m_ correlates with rheobase in Type II DGCs from E4-KI, but not E3-KI, mice; Pearson’s correlation analysis (two-sided). **n,** C_m_ correlates with spike latency in Type II DGCs from E4-KI, but not E3-KI, mice; Pearson’s correlation analysis (two-sided). All data are represented as mean ± s.e.m., *p<0.05, **p<0.01, ***p<0.001, ****p<0.0001, unpaired two-tailed t-test or Mann-Whitney U test. N= 4 and 3 for young E3-KI and E4-KI mice, respectively; N= 5 each for aged E3-KI and E4-KI mice.

**Extended Data Figure 4.**
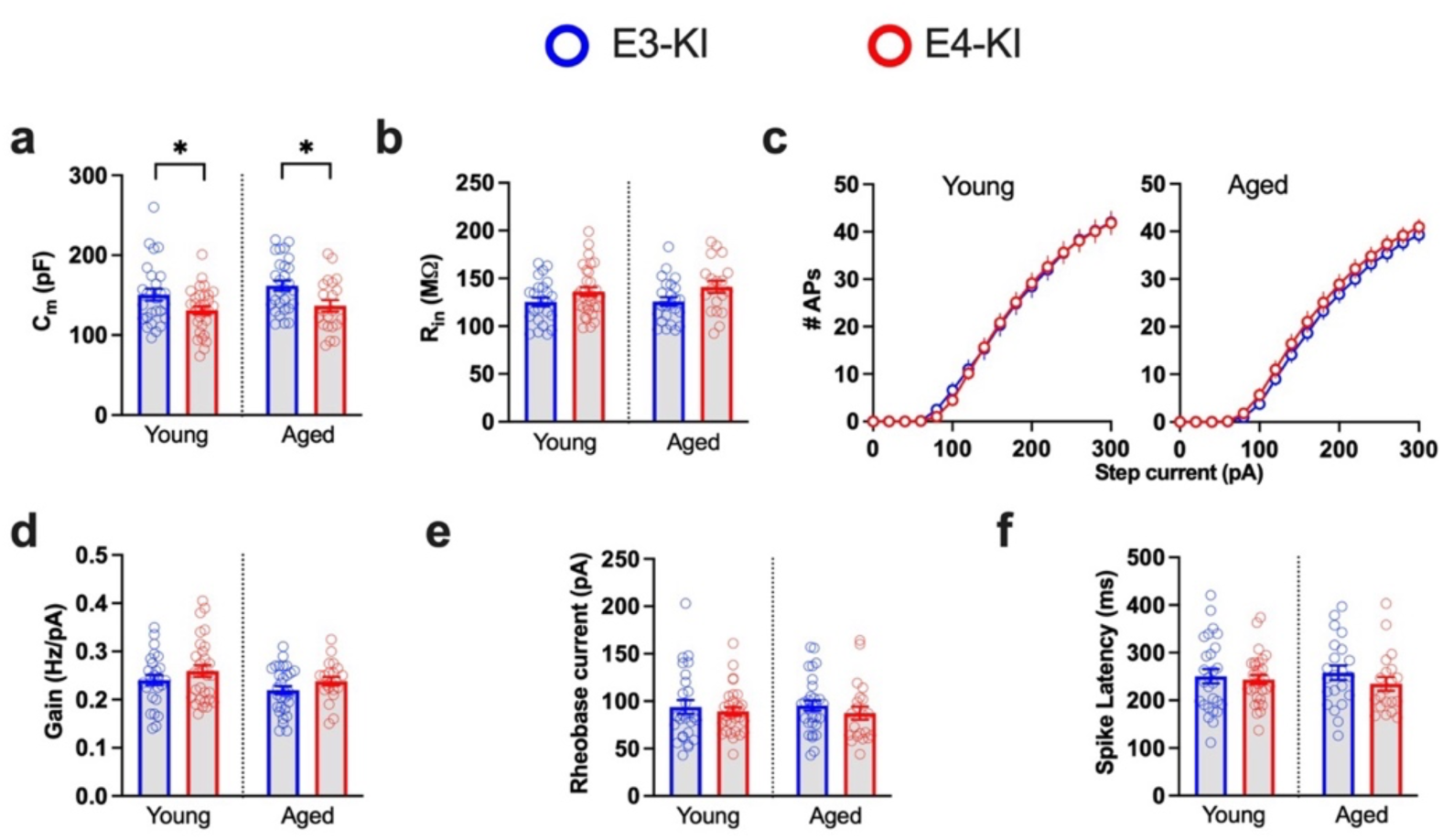
Lack of APOE4-driven hyperexcitability phenotype in hippocampal CA1 pyramidal cells (PCs). **a,** Cell capacitance (C_m_) values. **b,** Input resistance (R_in_) values. **c,** Input current – firing rate relationship (I-F Curve), **d,** Firing gain values. **e,** Rheobase values. **f,** Spike Latency values. All data are represented as mean ± s.e.m., *p<0.05, unpaired two-tailed t-test or Mann-Whitney test. N = 4 mice each for young E3-KI and E4-KI mice and 4–6 for aged E3-KI and E4-KI mice.

**Extended Data Figure 5.**
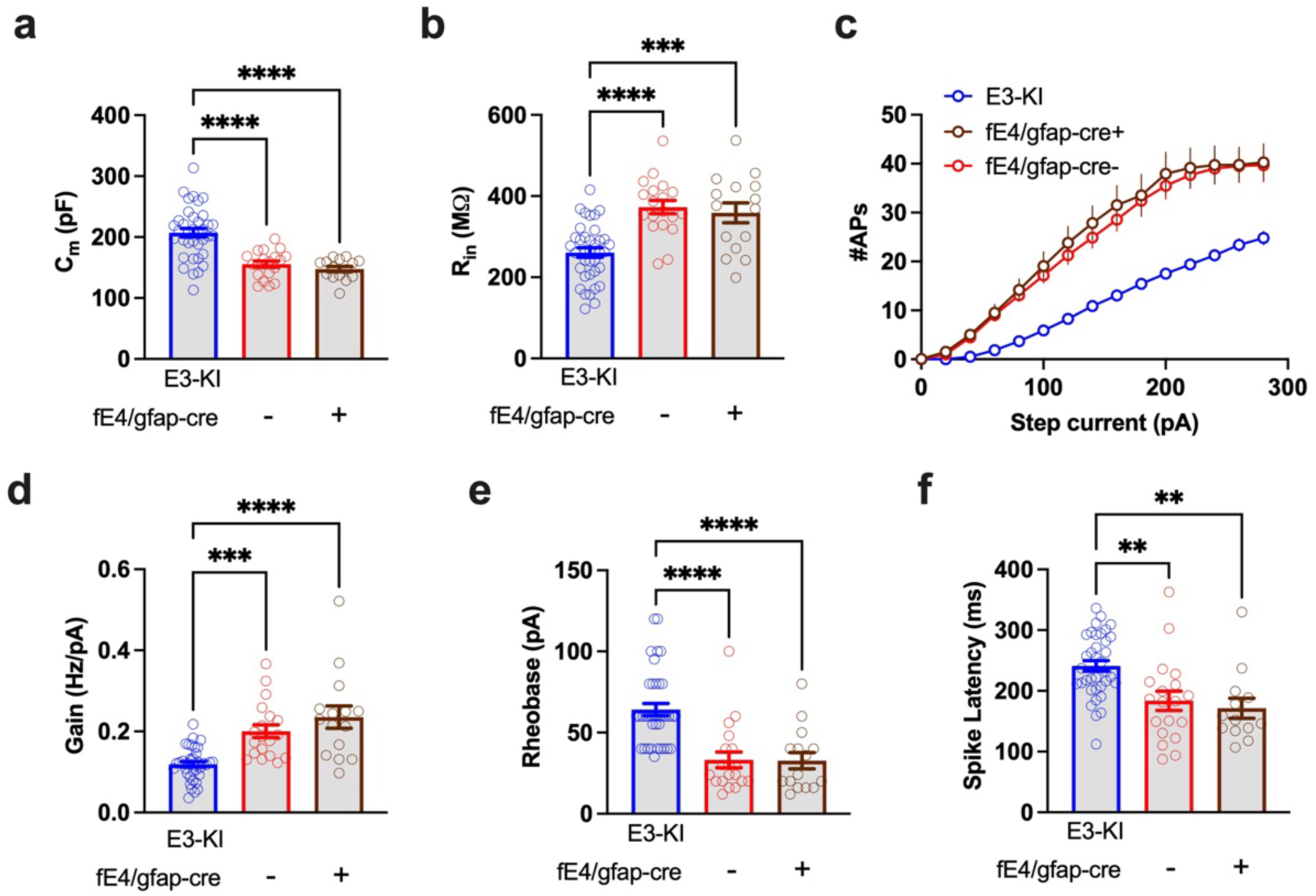
Selective APOE4 removal from astrocytes has no effect on the morpho-electric and excitability phenotypes of CA3 PCs in young fE4-KI^GFAP-Cre+^ mice. **a,** Cell capacitance (C_m_) values. **b,** Input resistance (R_in_) values. **c,** Input current – firing rate relationship (I-F Curves), **d,** Firing gain values. **e,** Rheobase values. **f,** Spike Latency values. All data are represented as mean ± s.e.m., **p<0.01,***p<0.001,****p<0.0001, one-way ANOVA with Tukey’s post-hoc multiple comparisons test. N= 8 or young E3-KI mice, and 3 for each fE4-KI^Syn1-Cre^ genotype.

**Extended Data Figure 6.**
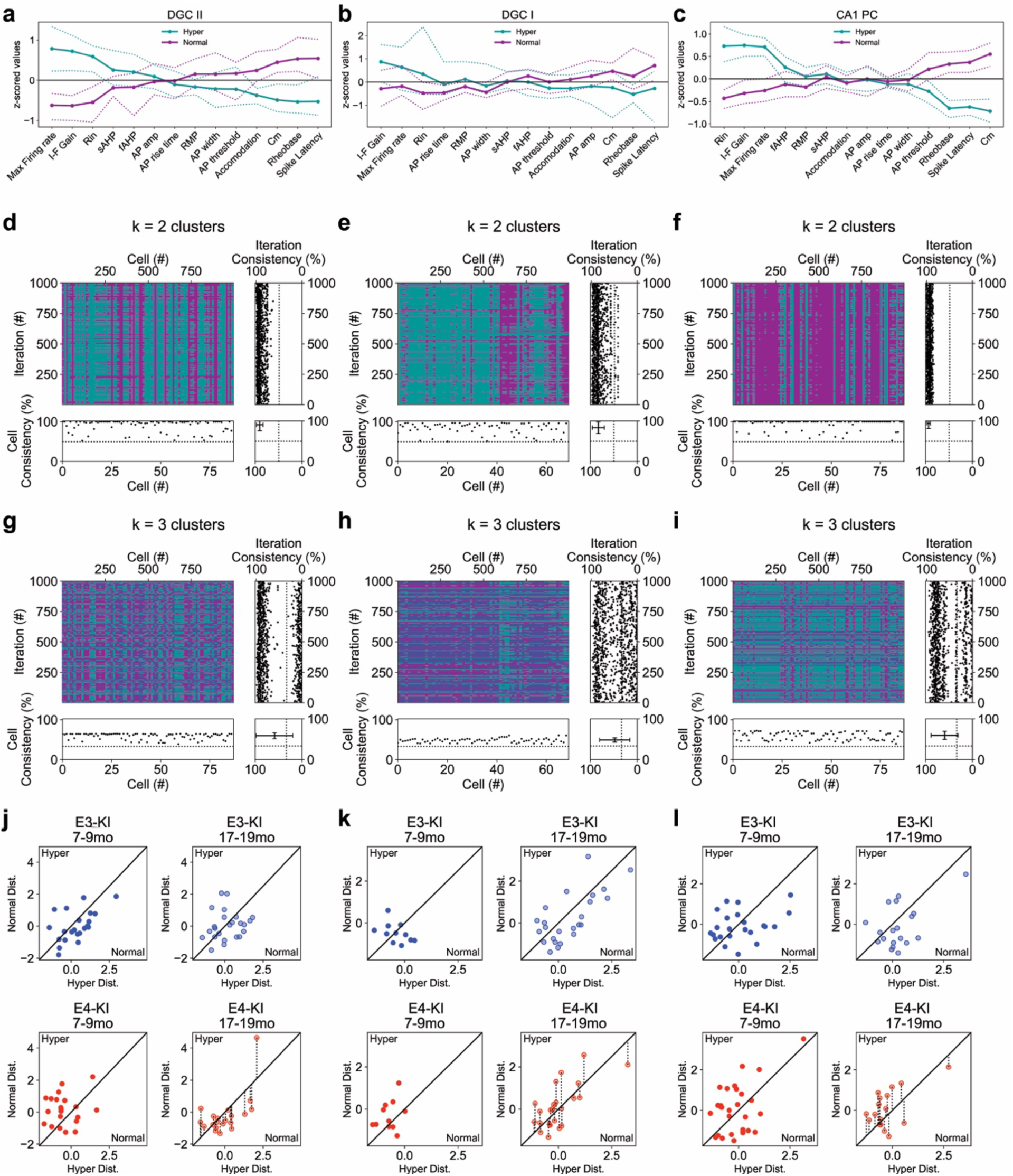
K-means clustering analysis of Type I DGC (DGC I), Type II DGC (DGC II), and CA1 PCs. **a–c,** Centroids determined by k-means clustering into two clusters for DGC II (a), DGC I (b), and CA1 PCs (c). Dashed lines indicate 99%CI determined from cross validation iterations. **d–f,** Cross validation analysis for with k=2 clusters for for DGC II (d), DGC I (e), and CA1 PCs (f). For each, heat map shows individual cell classification across each iteration. Upper Right) panel displays the probability of each iteration matching the classification of the full dataset. Bottom Left) panel displays the probability of a cell maintaining is classification identity across iterations. Lower Right) mean probabilities across iterations. Dashed lines indicate chance probability (50%). **g–i,** Cross validation analysis for with k=3 clusters for for DGC II (g), DGC I (h), and CA1 PCs (i). **j–l,** Scatter plots indicating the comparative distance from centroid for each cell binned by age and genotype for for DGC II (j), DGC I (k), and CA1 PCs (l). Unity line represents the decision line between each hyperexcitable and normal. ***p<0.001, ****p<0.0001, two-way ANOVA with Tukey’s post hoc multiple comparisons test.

**Extended Data Figure 7.**
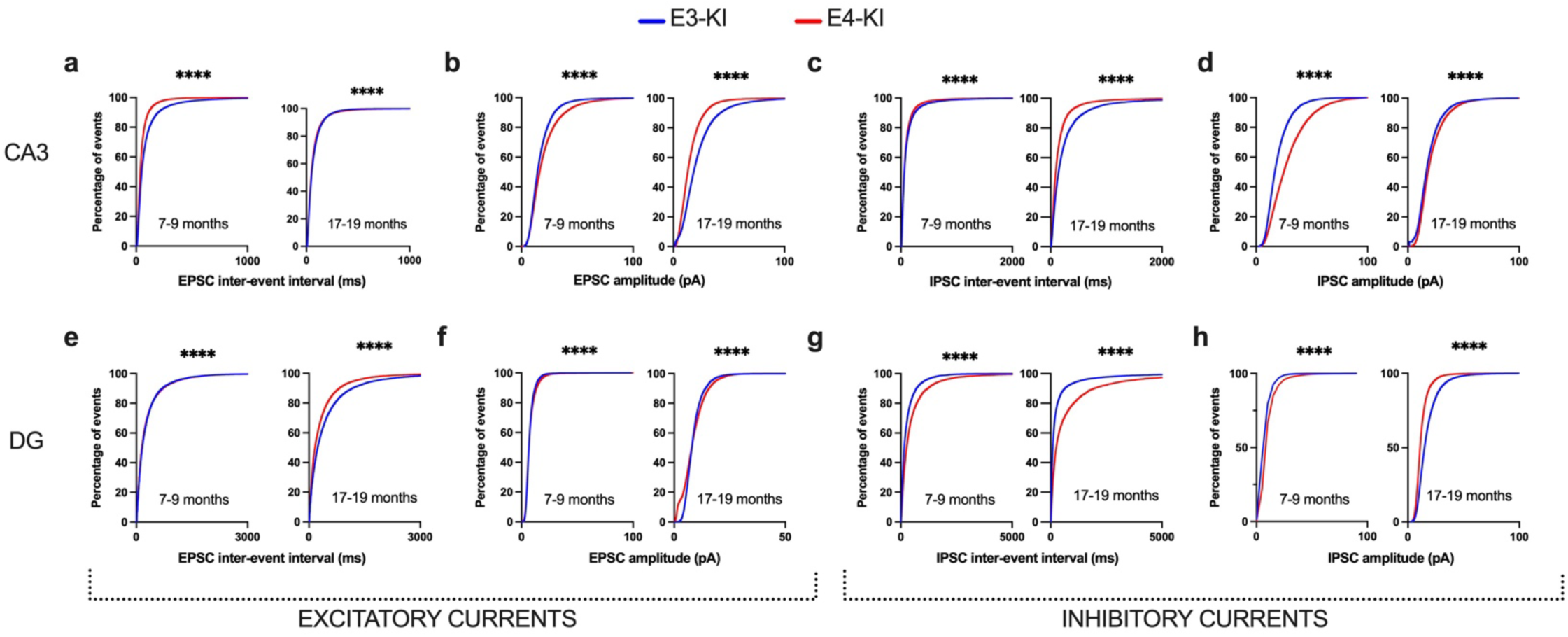
Cumulative distributions for inter-event intervals and amplitudes of all synaptic events shown in Figure 6. First 1000 events in each recording were used for the analysis. **a,** CA3 PC EPSC inter-event intervals. **b,** CA3 PC EPSC amplitudes. **c,** CA3 PC IPSC inter-event intervals. **d,** CA3 PC IPSC amplitudes. **e,** DGC EPSC inter-event intervals. **f,** DGC EPSC amplitudes. **g,** DGC IPSC inter-event intervals. **h,** DGC IPSC amplitudes. ****p<0.0001, Kolmogorov-Smirnov test.

**Extended Data Figure 8.**
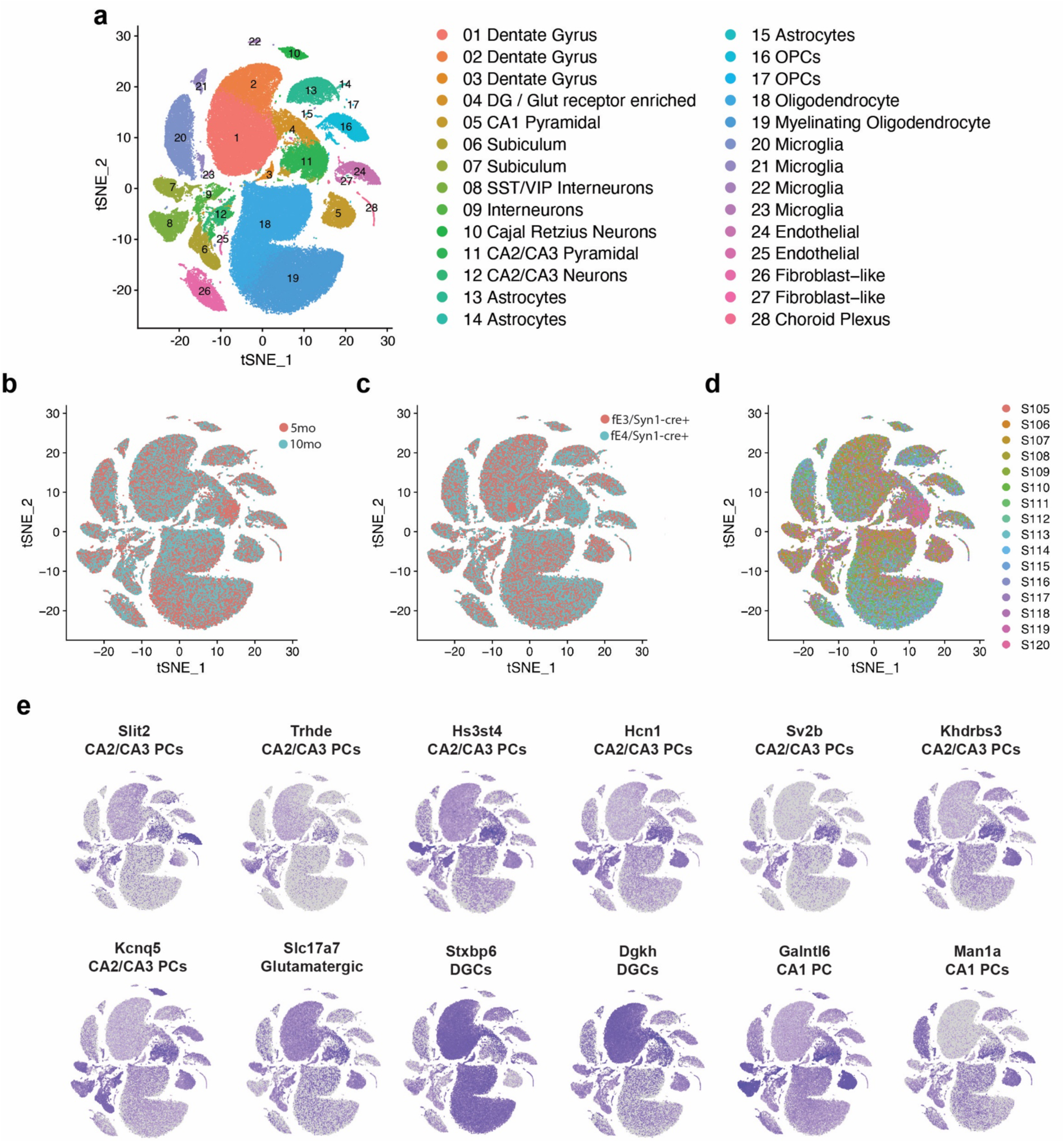
Generation of snRNA-seq datasets from the hippocampi of fE-KI^Syn1-Cre+^ mice for analysis of neuron-specific APOE effects. Hippocampal snRNA-seq data were collected from fE4^Syn1-Cre+^ and fE3^Syn1-Cre+^ mice at 5 and 10 months (N = 4 mice per genotype per age, total 16 mice). The hippocampi were dissociated, and nuclei were labeled with DAPI and isolated using flow cytometry before processing using the 10x Chromium v3 system**. a**, The snRNA-seq data were filtered, normalized, and clustered using the Seurat package, which yielded 28 distinct cellular clusters, including 12 neuronal clusters (Clusters 1–12) and 16 non-neuronal clusters (Clusters 13–28). Cell type identities for each cluster were identified using positively enriched marker genes. The tSNE plot shows numbered clusters; the corresponding key shows cell types for each numbered cluster. **b–d**, Single nuclei from both ages (b), genotypes (c) and individual mice (d) are distributed across all 28 clusters. **e,** Feature plots of marker genes for primary excitatory cell types.

**Extended Data Figure 9.**
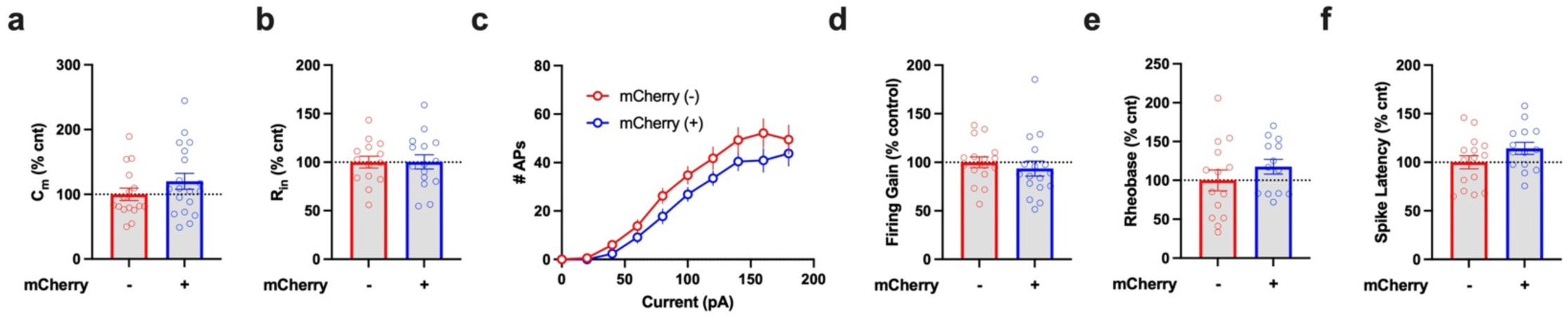
Lack of Mdga2 CRISPRi effects on neuronal morpho-electric and excitability parameters in young E4-KI mice. Mdga2 CRISPRi/mCherry+ DGCs display unaltered morpho-electric and excitability parameters: C_m_ (a), R_in_ (b), I-F Curves (c), firing gain (d), rheobase (e), and spike latency (f). All data are represented as mean ± s.e.m., differences examined using unpaired two-tailed t-test or Mann-Whitney U test; N= 5 for each condition.

